# Spatial signature of resource distribution is mediated by consumer body size and habitat preference

**DOI:** 10.1101/2024.06.24.600507

**Authors:** Chelsea J. Little, Pierre Etienne Banville, Adam T. Ford, Rachel M. Germain

## Abstract

Consumers shape spatial patterns on landscapes by amplifying or dampening environmental heterogeneity through feeding, excretion, and movement of resources. The degree to which the environment is modified by consumers depends on species’ traits, including body mass, movement and foraging behavior, and habitat specialization. Global change is altering the size and traits of consumer populations, but our understanding of how this may impact resource heterogeneity is limited. Here, we developed an individual-based model of habitat specialists’ and generalists’ movement and activity in a patchy landscape and investigated the impact of changes in population and mean body sizes on landscape-scale resource heterogeneity. We found that consumers specializing on low-resource habitats (a common risk avoidance strategy) increased spatial resource heterogeneity regardless of their population and body size. By contrast, generalists eroded differences among habitats, and we further found that resource heterogeneity decreased with the average body size of generalist consumers, even while controlling for total consumer biomass. Larger perceptual ranges increased the area over which generalist consumers could select foraging habitat, and reduced the extent to which they eroded landscape structure. These nuanced spatial outcomes of consumer-resource interactions emerge from how metabolic demands, which scale nonlinearly with body size, play out among habitat types which attract different consumers, as well as the scale at which those consumers make habitat selection decisions. Since global change disproportionately impacts larger species and specialists, indirect consequences on ecosystems may arise via biotic processes, affecting spatial heterogeneity of future landscapes.

## Introduction

Resource landscapes are often thought of as an environmental template that organisms respond to, but this neglects the role that organisms play in shaping spatial distributions of resources. Animal consumers serve as important resource vectors in landscapes (Earl and Zollner 2017, McInturf et al. 2019, Little et al. 2022, Pringle et al. 2023). The deposition of resources via feces, urine, seeds, placenta, and carcasses contribute to resource dispersal in the landscape (McIntyre et al. 2008, Bump et al. 2009, Sitters et al. 2017, Schmitz et al. 2018, le Roux et al. 2020, Ferraro et al. 2023). Organismal activity will therefore impact spatial resource heterogeneity, which in turn facilitates the coexistence of species (Amarasekare 2003) and thus ecosystem functioning (Loreau et al. 2003b). The distribution of resources in space is therefore an important landscape characteristic that shapes patterns of biodiversity and ecosystem functioning (Tylianakis et al. 2008, Seiferling et al. 2014, Gonzalez et al. 2020, Rizzuto et al. 2024). With global change altering abiotic conditions and consumer populations, it is imperative to gain a better understanding of the processes shaping spatial resource heterogeneity.

The traits of species making up consumer communities may provide a key to understanding organismal effects on resource landscapes. A handful of studies have established that species’ traits, such as behavior and body size, determine the amount, stoichiometry, and scale of consumer-driven resource cycling (Sitters et al. 2017, Veldhuis et al. 2018, Subalusky and Post 2019). For example, diet preferences and risk avoidance strategies may concentrate consumer activity into specific habitat types, leading to divergent rates of resource cycling across the landscape (Peller et al. 2022). Meanwhile, consumer body size influences resource dispersal through at least two pathways. First, energetic requirements scale nonlinearly with body size (Brown et al. 2004), and consequently so should resource consumption and excretion. Second, body size is associated with differences in movement ability and perceptual range (Little et al. 2022, Albaladejo-Robles et al. 2023, Straus et al. 2024), and thus the spatial dispersion of foraging and resource deposition.

It is challenging to predict how changing consumer communities in the Anthropocene will rewire resource distributions, because the same traits that mediate species’ responses to global change also determine their impacts on resource cycling. A major prediction of global change is the loss of large, specialist consumers, as increased metabolic requirements in warmer conditions and increased variability in resource supply through time increasingly causes resource demands to outpace supply (Clavel et al. 2011, Sheridan and Bickford 2011, Albaladejo-Robles et al. 2023, Martins et al. 2023). Concurrently, many mammalian species’ population sizes are declining in the Anthropocene (Finn et al. 2023). Examining the linkages between population density and resource spatial variation using caribou as a focal species, Ferraro *et al*. (2022) used an individual-based model to show that higher consumer densities more strongly exploited the resource landscape and increased spatial variation by creating resource hotspots (via carcass deposition) and coldspots (via resource consumption). This suggests that declining population densities could erode the mechanisms which create spatial variability in resource concentrations on landscapes. At even larger spatial scales, empirical work has shown that higher densities of bison alter nutrient cycling in ways that lead to higher-quality resources being available for longer periods of time in local areas of grassland landscapes; conversely, population decline alters the timing and quality of resources available to the entire ecosystem (Geremia et al. 2019). Over longer timescales, considering communities rather than single species, turnover may lead to more smaller, generalist consumers. The consequences of these shifts are more difficult to predict because body size and population density often trade off (Gjoni and Glazier 2020), and thus changes in body size might offset the effects of changes in density on resource distribution and lead to unique spatial patterns that could not be predicted from either dynamic alone. Simulations can provide a powerful tool for disentangling the individual and combined effects of population density, body size, and associated traits that affect resource use, such as species’ perceptual range and movement ability.

In this study, we present an empirically-calibrated, individual-based model of consumer movement where we factorially manipulated i) population density, ii) body size, iii) risk-based habitat preferences, and iv) perceptual range, and investigated effects on landscape-scale resource heterogeneity in a patchy landscape. Our model was parameterized to capture key biological features of consumers with two foraging strategies—generalists, and specialists that avoid high-nutrient resource patches (a common predator avoidance strategy)—where movement distances, metabolic requirements, and perceptual ranges are scaled with body size.

We hypothesized that specialists would accentuate the differences between nutrient-rich and nutrient-poor parts of the landscape by removing resources from lower-quality areas, and thus greater densities (compared to lower densities) of specialists would increase landscape-scale resource heterogeneity. By contrast, we hypothesized that greater densities (compared to lower densities) of generalists would preferentially graze high-quality areas until they resemble lower-quality areas, reducing spatial resource heterogeneity. To accompany this hypothesis, we further developed a suite of predictions about how consumer body size and total consumer biomass would shape spatial resource distributions (**Table 1**). Finally, we predicted that increasing the perceptual range and movement ability of consumers would reduce spatial variability of their contributions to nutrient dynamics.

**Table 1:**
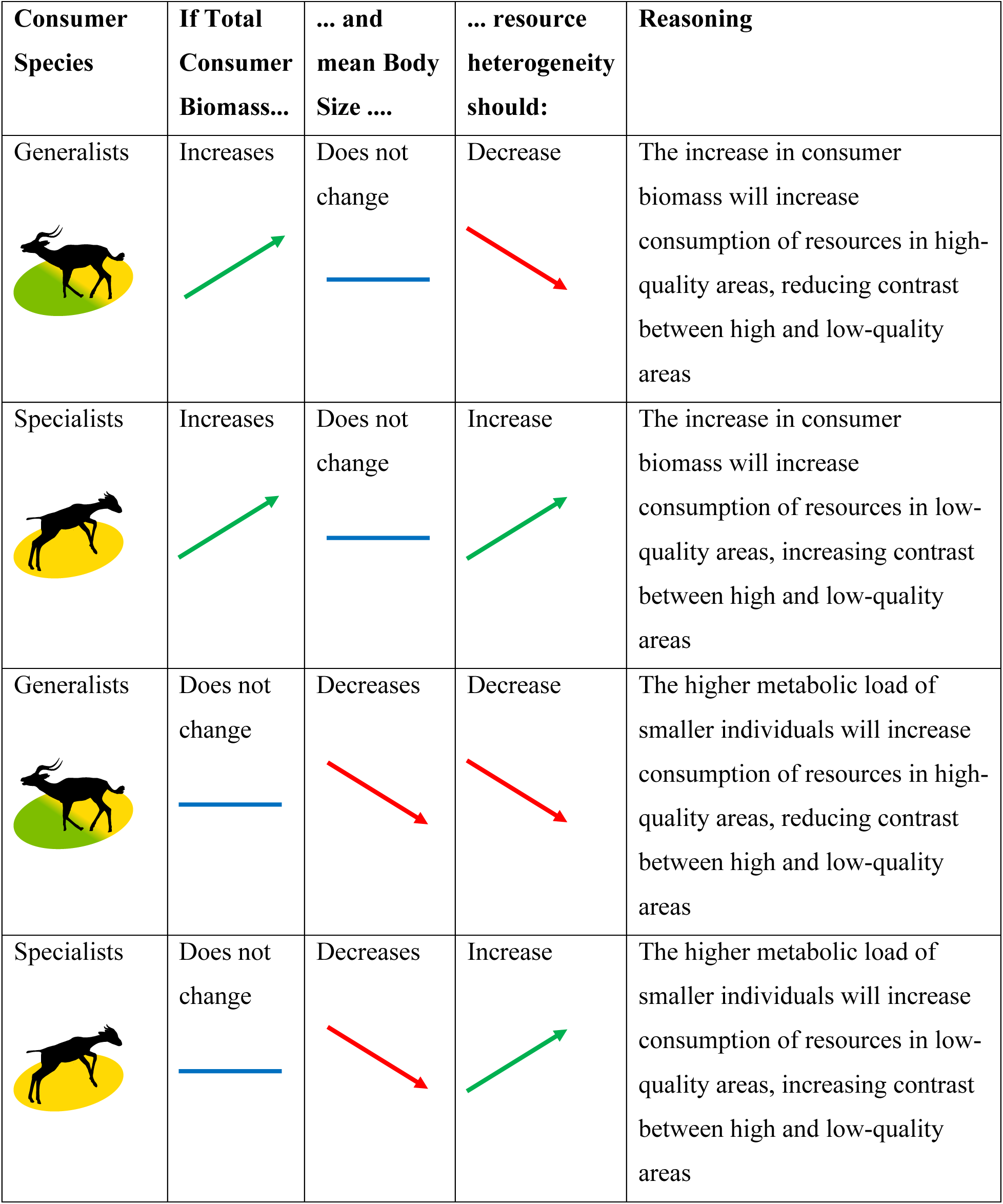
Hypothesized effect on resource heterogeneity of changing consumer species, total consumer biomass and mean body size on the landscape.

| Consumer Species | If Total Consumer Biomass... | ... and mean Body Size .... | ... resource heterogeneity should: | Reasoning |
| --- | --- | --- | --- | --- |
| Generalists | Increases | Does not change | Decrease | The increase in consumer biomass will increase consumption of resources in high-quality areas, reducing contrast between high and low-quality areas |
| Specialists | Increases | Does not change | Increase | The increase in consumer biomass will increase consumption of resources in low-quality areas, increasing contrast between high and low-quality areas |
| Generalists | Does not change | Decreases | Decrease | The higher metabolic load of smaller individuals will increase consumption of resources in high-quality areas, reducing contrast between high and low-quality areas |
| Specialists | Does not change | Decreases | Increase | The higher metabolic load of smaller individuals will increase consumption of resources in low-quality areas, increasing contrast between high and low-quality areas |

## Methods

### Modeling approach

We used NetLogo v6.0.4 (Wilensky 1999) to model generalists and specialists moving in a landscape, where consumers withdraw resources by foraging and deposit resources through feces and carcasses. This individual-based modeling framework has been used to effectively explore foraging movement and animal-transported subsidies in several past applications (e.g. Buchmann et al. 2012, Teckentrup et al. 2018, Bampoh et al. 2019, Ferraro et al. 2022). Key parameters are indicated in **Table 2**, and a detailed explanation is found in the model Overview, Design, and Details (ODD) (Grimm et al. 2010) in Appendix I. This landscape is divided into a 141 x 141 grid of equal-sized pixels, each pixel representing 100 m^2^. The landscape has two discrete habitat types (**Fig. 1**), approximating a Kenyan savanna (see Ford et al. [2014]): open, high-nutrient glades as clusters of pixels (each cluster forming a “Glade”), in a “Matrix” of bushy, lower-quality grassland (Augustine 2004). Each pixel is classified as Glade or Matrix and is assigned a forage amount drawn from a normal distribution centered on 50 g/m^2^ for Glades and 33 g/m^2^ for Matrix, with a standard distribution of 25% of the mean; forage contained 6% nitrogen (N) in Glades and 4% in Matrix. Landscapes contained an average of four Glades, making up ∼5% of the landscape area (**Fig. 1**).

**Figure 1.**
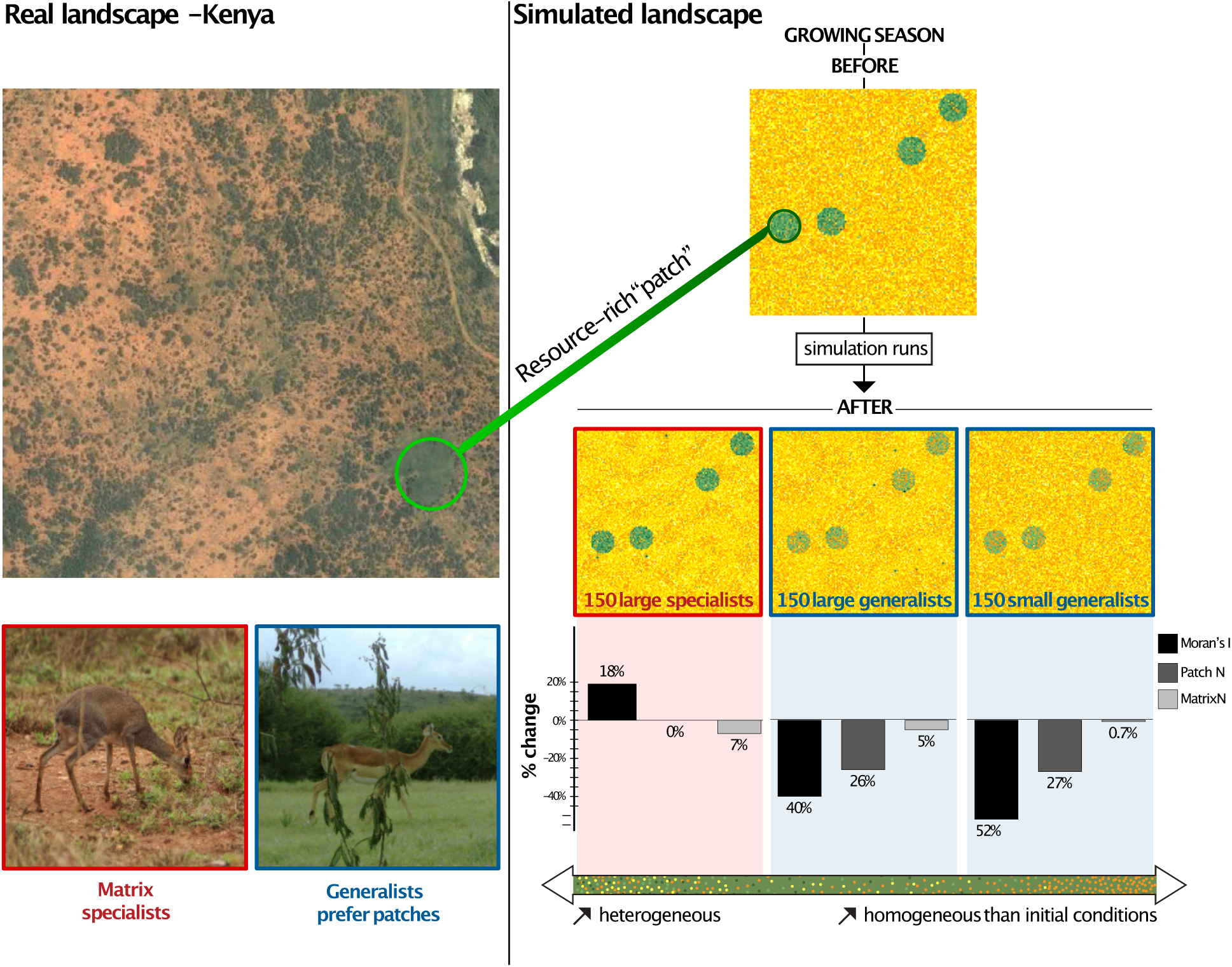
N distributions within Glades and Matrix at the beginning (top right) and end (middle right) of the growing season in simulated landscapes, which are based on a Kenyan savanna (left) with habitat specialists that forage only in Matrix, and generalists that forage across the whole landscape but prefer Glades. In our model, consumers realistically increase perceptual range, movement ability, and metabolic needs with body size. Pixel-level N concentration ranges from low (yellow) to intermediate (orange) and high (green). In the middle right, N distributions after grazing, movement, and N deposition for three example scenarios out of the study’s total 64 (see Table S1): here, by equal numbers of large (40kg) specialists, large generalists, or small (10kg) generalists in example simulations. Percent changes in Moran’s I, a measure of spatial autocorrelation (Moran 1950), and in the nitrogen content of each pixel in Glades and Matrix are indicated for each simulation in the bottom right panels. In the “150 large specialists” panel, note the waves of N distribution in the Matrix created by foraging specialists compared to the initial landscape. Small dark green spots at the end of the growing season represent N deposition by carcasses of consumers that died during the simulation, which is most evident for larger consumers. Real landscape image © 2024 Maxar Technologies.

**Table 2:** Key parameters influencing the initial distribution of N on the landscape and the withdrawals and deposits of N by animals.

| Parameter (units) | Value | Reference |
| --- | --- | --- |
| Mean forage per Glade pixel (g) | 5000 | Augustine 2004 |
| Mean forage per Matrix pixel (g) | 3300 | Augustine 2004 |
| Mean N per Glade pixel (g) | 300 (6% of forage) | Augustine 2004 |
| Mean N per Matrix pixel (g) | 132 (4% of forage) | Augustine 2004 |
| N in consumers' body (g) | 2% * bodymass | Pace and Rathbun 1945 |
| Daily metabolic needs of consumers for forage (g) | $(7.94 * (\text{bodymass})^{0.646} / 11.5)$ | Nagy 2001 |
| Yearly % chance of death for animals | 10% for specialists; 10% in Glade, 13% in Matrix for generalists | Ogutu et al. 2012, Owen-Smith and Goodall 2014 |
| N deposited on landscape per defecation event (g) | 50% of N consumed since last defecation event | Singer and Schoenecker 2003 |
| N deposited on landscape per death event (g) | All body N + 100% N consumed since last defecation event | For spatial distribution of carcass N to pixels, see Appendix I |

Individuals in the model are either generalists or specialists. Specialist consumers, resembling Guenther’s dik-diks, stay in the Matrix to avoid predators (Kingswood and Kumamoto 1996, Otieno et al. 2019), while generalists, akin to impalas, can use both habitat types (Ford et al. 2014, Otieno et al. 2019). In nature, impalas are larger than dik-diks, however in our simulations we factorially manipulated habitat specialization and body size to also investigate landscape impacts of large specialists and small generalists, the latter of which may be predicted to become more common in the Anthropocene through community turnover. In our model, animals forage based on their metabolic needs and move within their perceptual range, both of which vary as a function of their body size (Nagy 2001, Mech and Zollner 2002, Forero-Medina and Vieira 2009, Hillaert et al. 2018, Straus et al. 2024). The association between body size and perceptual range is rarely tested empirically (but see previous references), therefore we chose three different parameterizations of the scaling parameter for sensitivity testing (see *Simulation Scenarios* below and the ODD in Appendix I). Foraging behavior adheres to the marginal value theorem, with animals selecting pixels with better-than-average forage available (Charnov 1976). Glade pixels are not considered by specialists, while generalists can choose any pixel (Glade or Matrix) and preferentially graze Glades at the start of each simulation because of the higher forage and N concentration. Mortality rate depends on species type (generalist or specialist) and location, with the two species types having different mortality rates in the Matrix (**Table 2**); this reflects that even at equal body sizes, other species traits such as behavioral risk avoidance shape predation risk (see Appendix I for details on movement, foraging, excretion, and death). Model timesteps represent 3 hours of movement/activity, and simulations were run for three months, or the length of one growing season. This time scale was chosen to investigate how animal foraging decisions in a period of resource abundance can shape the environmental template for consumer-resource interactions in future seasons and years.

### Simulation Scenarios

We designed our scenarios (**Table S1**) to look at the impact of 4 different factors on landscape heterogeneity: species role (habitat generalists or specialists), body size, perceptual range, and total biomass on the landscape, where the total biomass on the landscape is determined by the mean body size multiplied by the number of individuals. We tested combinations of body size (10kg, 25kg or 40kg), and total biomass on the landscape (varying between 500kg and 6000kg) for each of the species. We varied the total biomass to allow for comparisons where the total number of individuals is the same (50, 100, or 150 individuals representing 25, 50, or 75 individuals/km^2^; for reference, population density estimates for impala are ∼20/km^2^ (Augustine 2004) and for dik-diks range up to 112/km^2^ (Komers and Brotherton 1997)) and where the total biomass is the same (1000kg, 2000kg, or 4000kg). Initially holding perceptual range constant, we varied those factors to test all combinations possible, resulting in 34 scenarios as described in **Table S1**.

We then ran each scenario with three different parameterizations of perceptual range, defined by a relationship between consumer body size and the range’s radius (Straus et al. 2024). We chose to model a linear relationship between body size and perceptual distance, as in Hillaert et al. (2018). As a result, the area of a consumer’s perceptual range (in km^2^) scaled quadratically with consumer body mass, according to the equation

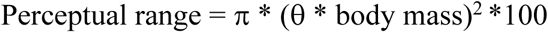

where θ was 0.25, 0.5, and 1 in our low-, medium- and high-perceptual range scenarios (see ODD in Appendix I for more details). Testing these different parameterizations of perceptual range added 30 more scenarios for a total of 64. This combination of body size and perceptual range scaling parameterization allowed us to effectively explore the outcomes of sensing behavior at different scales on the landscape. With the smallest body size and low perceptual range, consumers could sense only the closest pixels to their own (pixel radius = 2.5), while with the largest body size and high perceptual range, their perceptual range covered half a square kilometer, or roughly one third of the entire experimental landscape.

All scenarios were replicated 25 times, with different random configurations of Glades in each replicate simulation. The Netlogo simulations were processed through R version 4.2.2 using the package ‘nlrx’ (Salecker *et al*. 2019). They were performed on the high-performance computer Cedar from Digital Research Alliance of Canada. We conducted additional sensitivity testing by varying the parameterization of mortality rate, N deposition frequency, and the proportion of consumed N that was deposited by excretion/egestion (**Fig. S2, Fig. S3**).

The outputs from the simulation experiments focus on changes that happen in the distribution of N on the landscape and are recorded at the pixel level. They include the initial and final forage amount, the initial and final N amount, the N eaten through foraging, the N deposited through defecation or animal death, the coordinates and whether the pixel is Glade or Matrix. We also recorded the number of times a pixel was grazed, defecated on, and the number of deaths that occurred on a given pixel to help with results interpretation.

### Model Analysis

We analyzed changes in the spatial and total N distribution across the landscape to determine the effect of species role, body size, population density, total biomass, and perceptual range on landscape heterogeneity. The key variables of interest were the initial and final N amount of each pixel. For each individual simulation, we calculated the four statistical moments (i.e. mean, variance, skewness and kurtosis) for the initial and final N amount across all pixels. Comparing the value of those statistics for the initial N and final N amount indicates the change in the total N distribution across the landscape resulting from the simulation. For each scenario, we then calculated the mean and standard deviation of the 25 replicates for all statistics calculated at the individual simulation level. For a given statistic, we then plotted the mean with a standard deviation error bar of the specific scenarios that we wanted to compare, such as different body sizes or different number of individuals on the landscape. All statistical analyses were performed using R version 4.2.2 and the skewness and kurtosis were calculated using the package ‘geodiv’ (Smith et al. 2021).

We also explored the spatial changes in N distribution by calculating the Global Moran’s I (“Moran’s I”) on initial and final N for each individual simulation. The Moran’s I is a statistical test that measures spatial autocorrelation, indicating how similar are the pixels from their neighbors (Moran 1950). The Moran’s I sums covariances between pixels and, in its standard form using Queen connectivity (pixels are neighbors if they share a border or a point), gives equal weight to the 8 closest neighbors (first order neighbors) of a given pixel in the calculations. An increase in Moran’s I means that the pixels are more autocorrelated, indicating spatial clustering of resources and increased spatial heterogeneity. A decrease in Moran’s I means that the pixels are less autocorrelated and the distribution of N is less clustered on the landscape.

To determine if the relative impact of body size on spatial autocorrelation varies depending on the size of the neighborhood considered, we also calculated Moran’s I using 4^th^ order neighbors (80 neighbors), 10^th^ order neighbors (440 neighbors) and 17^th^ order neighbors (1224 neighbors) with an inverse distance spatial weight (i.e. the correlation between the closest neighbors weighs more than the correlation between neighbors further away). The 4^th^, 10^th^, and 17^th^ order neighbors were selected to approximate the perceptual range of individuals with a body size of 10kg, 25kg, and 40kg respectively. Individuals with larger body sizes can access pixels that are located further away than individuals with smaller body sizes, but are more likely to select nearby pixels to minimize energy costs. We found no qualitative differences in the results between the “queen” configuration Moran’s I and Moran’s I with 4^th^, 10^th^, and 17^th^ order neighbors, and hence present results with the standard Moran’s I only. The Moran’s I analysis was conducted using the package ‘spdep’ (Bivand and Wong 2018) in R version 4.2.2.

To better understand the drivers of the changes in the spatial and total N distribution at the landscape level, we analyzed the variables making up the final amount of N (for a given pixel, Final N = Initial N + N Deposited – N Eaten) at the landscape level and separately for Glades and Matrix. We calculated the mean and variance for N deposited, N eaten, and net N (difference between N Deposited and N Eaten) for the landscape as a whole and for Glades and Matrix separately for each individual simulation. As potential explanatory variables for N dynamics, we calculated the mean and variance in the number of times a pixel was grazed and defecated on, and the total number of death events at the landscape level for each individual simulation.

## Results

Spatial distribution of N varied across simulated landscapes at the end of the growing season. Although larger body sizes and larger population biomass led to more forage consumed (**Fig. 2**) – as expected based on the metabolic scaling built into our model – and greater N depletion (**Fig. S1**) in the total landscape for both specialists and generalists, changes in the landscape-scale spatial heterogeneity of N resulted from the interplay between N depletion in Glades and Matrix. Increasing or decreasing consumers’ mortality rate had minimal effects on total forage consumption (**Fig. S2**). Below, we describe how species’ traits and population densities shaped resource distributions across simulated landscapes.

**Figure 2.**
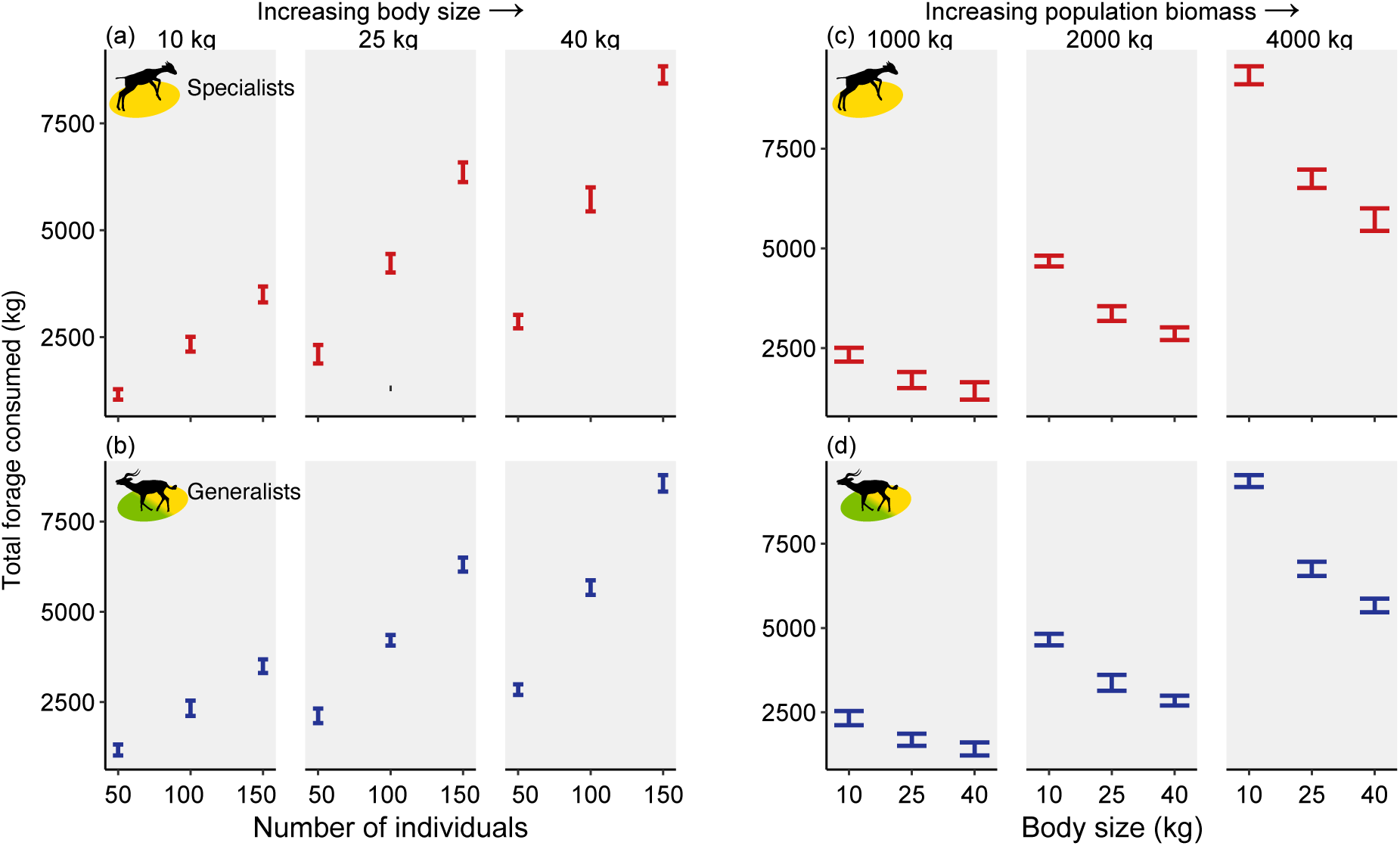
Total forage consumed during simulations with different combinations of parameters. (a) and (b) show the effect of increasing population density (on the x-axis) at small, medium, or large mean individual body size (left, center, and right panels), when the landscape is populated either with specialists that forage only in Matrix (a) or generalists that forage in both habitats (b). (c) and (d) show the effect of increasing mean body size (on the x-axis) when the total population biomass on the landscape is 1000 kg (left column), 2000 kg (middle column), or 4000 kg (right column) of specialists (c) or generalists (d). Bars show ± one standard deviation of 25 simulations for each scenario.

### Specialists increase (and generalists decrease) resource heterogeneity

As predicted, habitat specialists foraging only in the Matrix accentuated differences between habitat types (note increase in Moran’s I, represented as positive values on the y axis, in **Fig. 3**, top panels); for these specialists, greater population density and, when holding population size constant, larger body sizes were associated with greater increases in spatial resource heterogeneity (**Fig. 3a**). However, when holding population biomass constant, larger body sizes were associated with smaller increases in spatial resource heterogeneity (**Fig. 3c**). Also as predicted, generalists, who preferentially forage in Glades due to their higher N content, homogenized the landscape, reducing differences between Glades and Matrix (note decrease in Moran’s I, represented as negative values on the y axis, in **Fig. 3**, bottom panels).

**Figure 3.**
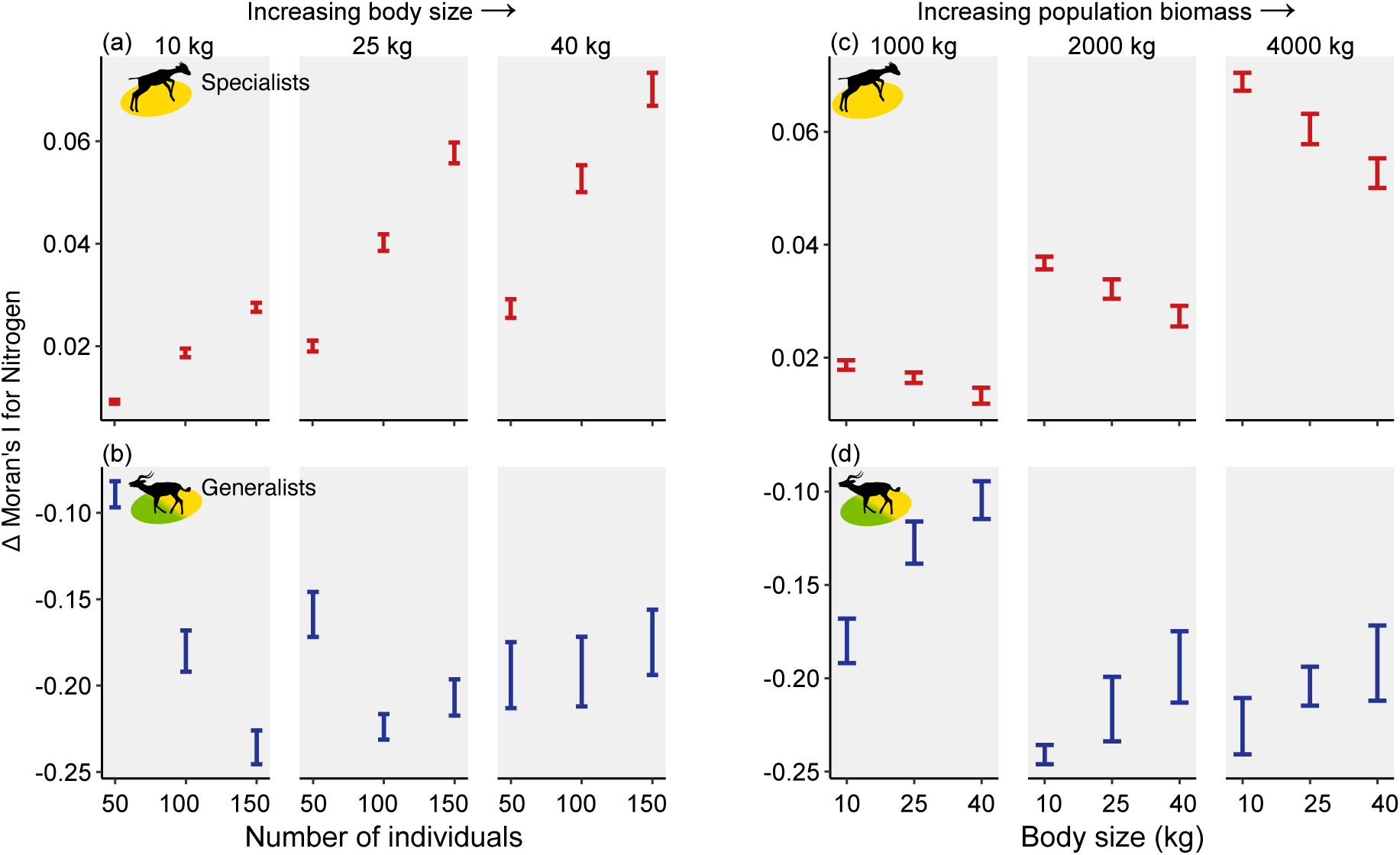
Change (Δ) in Moran’s I from the beginning and end of simulations for the spatial distribution of Nitrogen. (a) and (b) show the effect of increasing population density (on the x-axis) at small, medium, or large mean individual body size (left, center, and right panels), when the landscape is populated either with specialists that forage only in Matrix (a) or generalists that forage in both habitats (b). (c) and (d) show the effect of increasing mean body size (on the x-axis) when the total population biomass on the landscape is 1000 kg (left column), 2000 kg (middle column), or 4000 kg (right column) of specialists (c) or generalists (d). Bars show ± one standard deviation of 25 simulations for each scenario. Note different y-axes in top and bottom panels.

### Population density effects on resource heterogeneity by small but not large generalists

For small generalists, higher population densities led to greater homogenization of N distribution over the course of the simulations than that caused by less dense populations (**Fig. 3**). Since small generalists, even at higher population densities, did not remove enough forage from the Glades to make them of comparable quality as the Matrix (**Fig. S3**), N removal from the Matrix was negligible (i.e., the first panel of **Fig. 4c**) and larger population densities were nearly linearly associated with reductions in Moran’s I (i.e., the first panel in **Fig. 4b**, and first panel of **Fig. 3b**).

**Figure 4.**
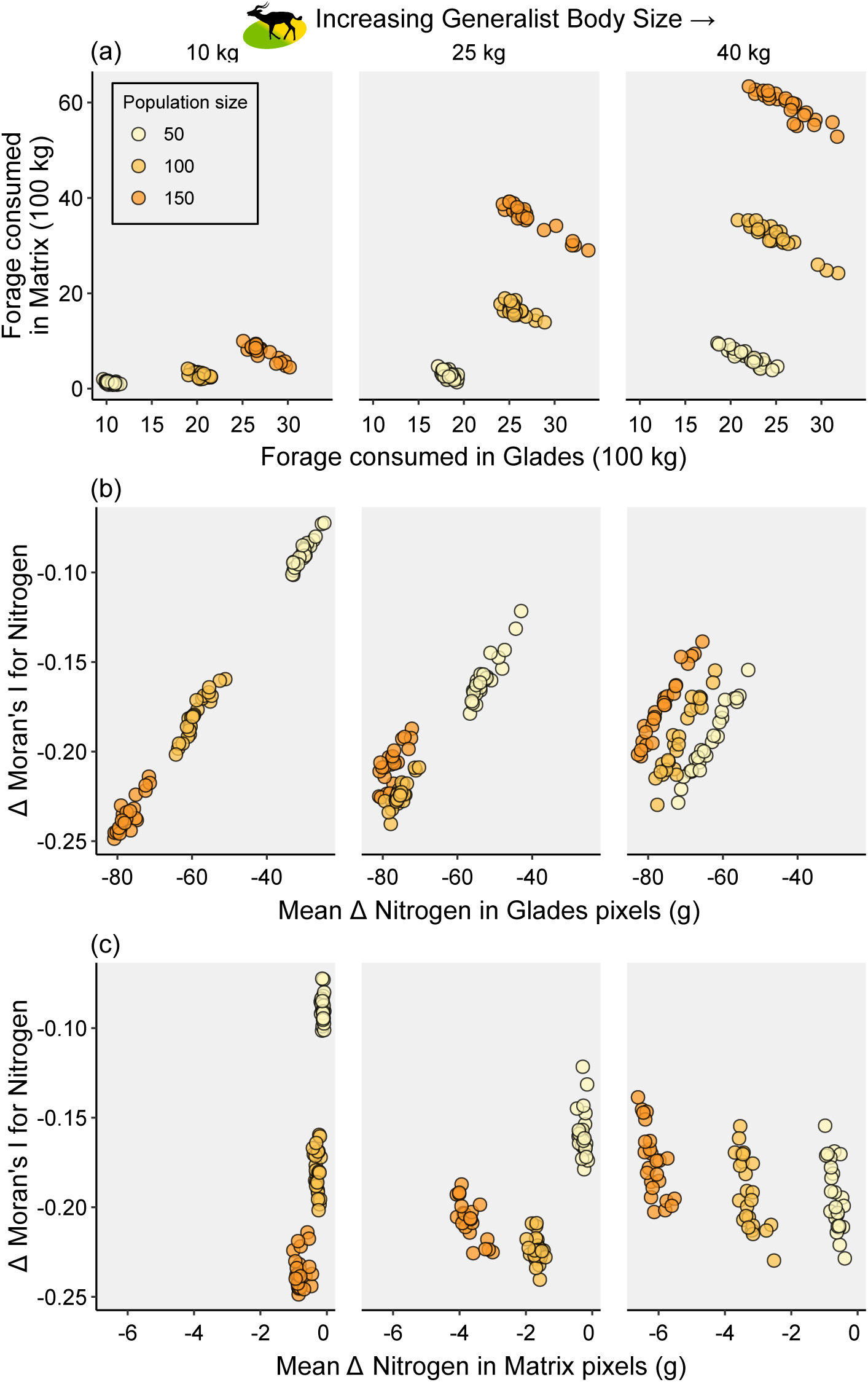
Changes in forage consumed in Glades and Matrix and in the spatial distribution of nitrogen in simulations with varying generalist body size and population density. Each point represents one simulation. (a) Relationship between the total forage consumed in Glades and the total forage consumed in the Matrix. Lower panels show change in Moran’s I with (b) net nitrogen loss from Glades and (c) net nitrogen loss from the Matrix.

However, for large generalists, the reduction in Moran’s I was fairly consistent across population densities, with slightly smaller reductions at higher densities. Larger generalists depleted Glades faster than smaller generalists, subsequently removing N from the Matrix (**Fig. 4a**) and probably occasionally returning to the Glades as the Matrix also became depleted. This led to similar reductions in spatial structure (measured by Moran’s I) across population densities (**Fig. 3b**), even if more forage was removed overall by larger populations (**Fig. 2b**). However, the increased N depletion of Glades by the highest densities of large generalists (x-axis in rightmost panel of **Fig. 4b**) was offset by their increased removal of N from the Matrix (x-axis in rightmost panel of **Fig. 4c**). As a result, the difference between Glade and Matrix quality (and thus Moran’s I) was comparatively stable across densities of large generalists (**Fig. 3b**), in contrast to variations caused by different densities of small generalists.

### Metabolic signature of body size is most apparent at low population densities

Comparing scenarios with equal total population biomass, but with many small versus fewer large consumers, revealed the effects of nonlinear scaling of metabolic requirements on spatial distribution of N (**Fig. 5**). As predicted by metabolic theory, total forage consumption was higher with many small consumers compared to few large ones (**Fig. 2**).

**Figure 5.**
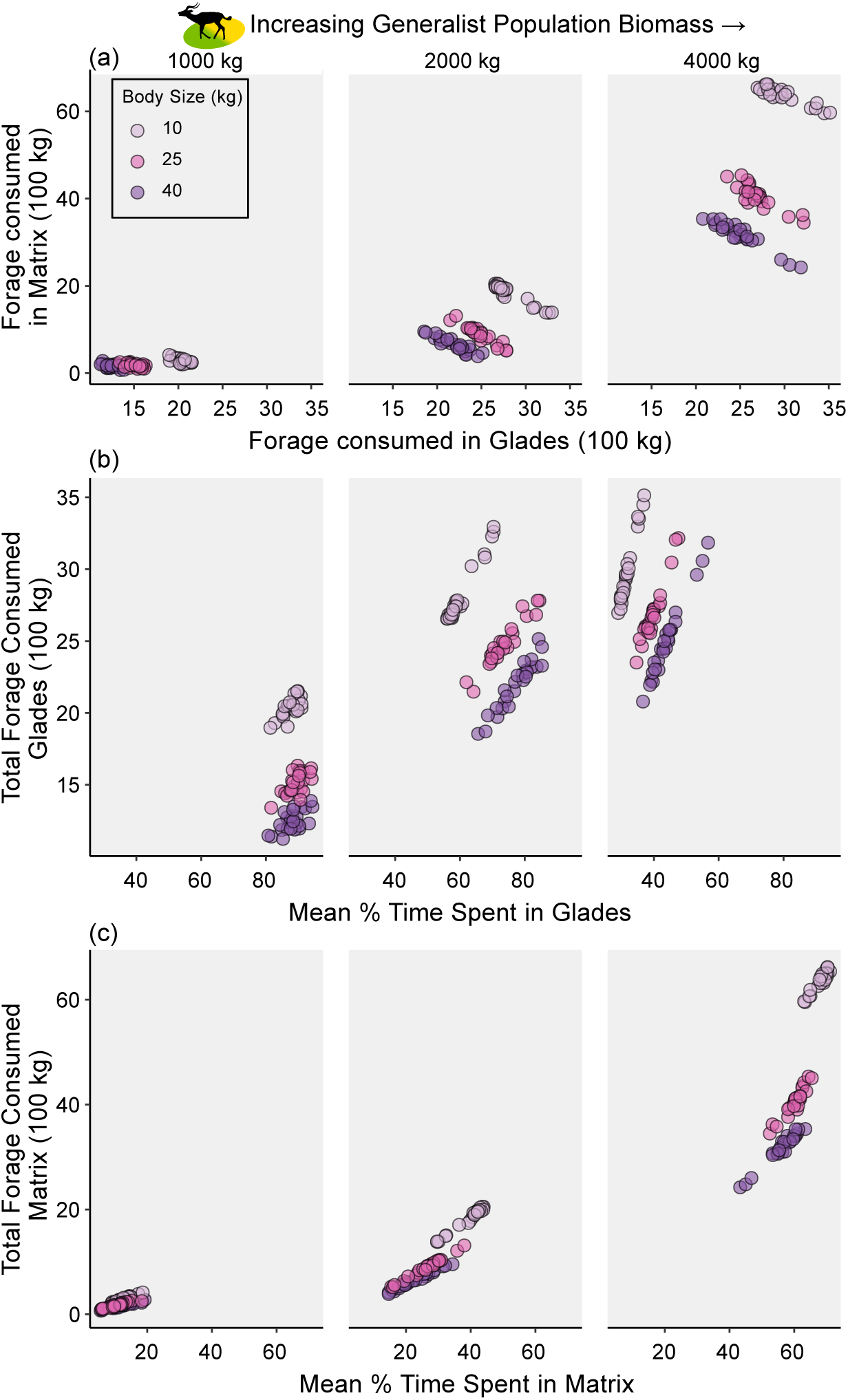
Changes in forage consumed in Glades and Matrix with different amounts of total generalist biomass on the landscape (shown in left, center, and right panels), and when that biomass is made up of different average body sizes (indicated by color of points). Each point represents one simulation. (a) Relationship between the total forage consumed in Glades and total forage consumed in the Matrix. (b) Relationship between the mean percentage of time individuals spent in Glades and the total forage consumed in Glades. (c) Relationship between the mean percentage of time individuals spent in the Matrix and total forage consumed in the Matrix.

At lower total biomass, generalists of all body sizes spent most of their time in Glades (**Fig. 5b)**. However, small generalists consumed more forage there (**Fig. 5b**), leading to a stronger homogenizing effect than their larger counterparts (**Fig. 3c**), as hypothesized.

On the other hand, at larger total biomass, small generalists grazed the Glades more thoroughly than large generalists (**Fig. 5a**) despite spending less time there (**Fig. 5b**), and instead spent upwards of 60% of their time in the Matrix (**Fig. 5c**). As a result, small generalists consumed markedly more forage in the Matrix than populations of large generalists (**Figs. 5a**, **5c**). Because this increased N removal from the Matrix made it even more resource-poor, this foraging behavior by high densities of small generalists thus maintained differences in N concentrations between Glades and Matrix pixels even if each respectively lost N in absolute terms. This is reflected in the weaker differences in resource spatial heterogeneity (measured by Moran’s I) arising from scenarios with dense populations of small vs. large generalists, compared to larger differences in Moran’s I arising from scenarios with sparse populations of small vs. large generalists (**Fig. 3d**).

### Increasing perceptual range of generalists maintains resource spatial heterogeneity

Increasing consumers’ perceptual ranges tended to alter foraging behavior, whereby generalists with larger perceptual ranges spent less time in Glades (**Fig. 6a**), with some exceptions. At intermediate and higher population biomasses, we found that as perceptual range increased, the net loss of nitrogen from Glades generally decreased, with the strongest trend occurring at the largest body sizes (**Fig. 6b**). Correspondingly, net loss of nitrogen from the Matrix increased slightly with increasing generalist perceptual range (**Fig. 6c**). As a result, spatial autocorrelation of N (measured by Moran’s I) remained higher when generalists had larger perceptual ranges (**Fig. 6d**).

**Figure 6.**
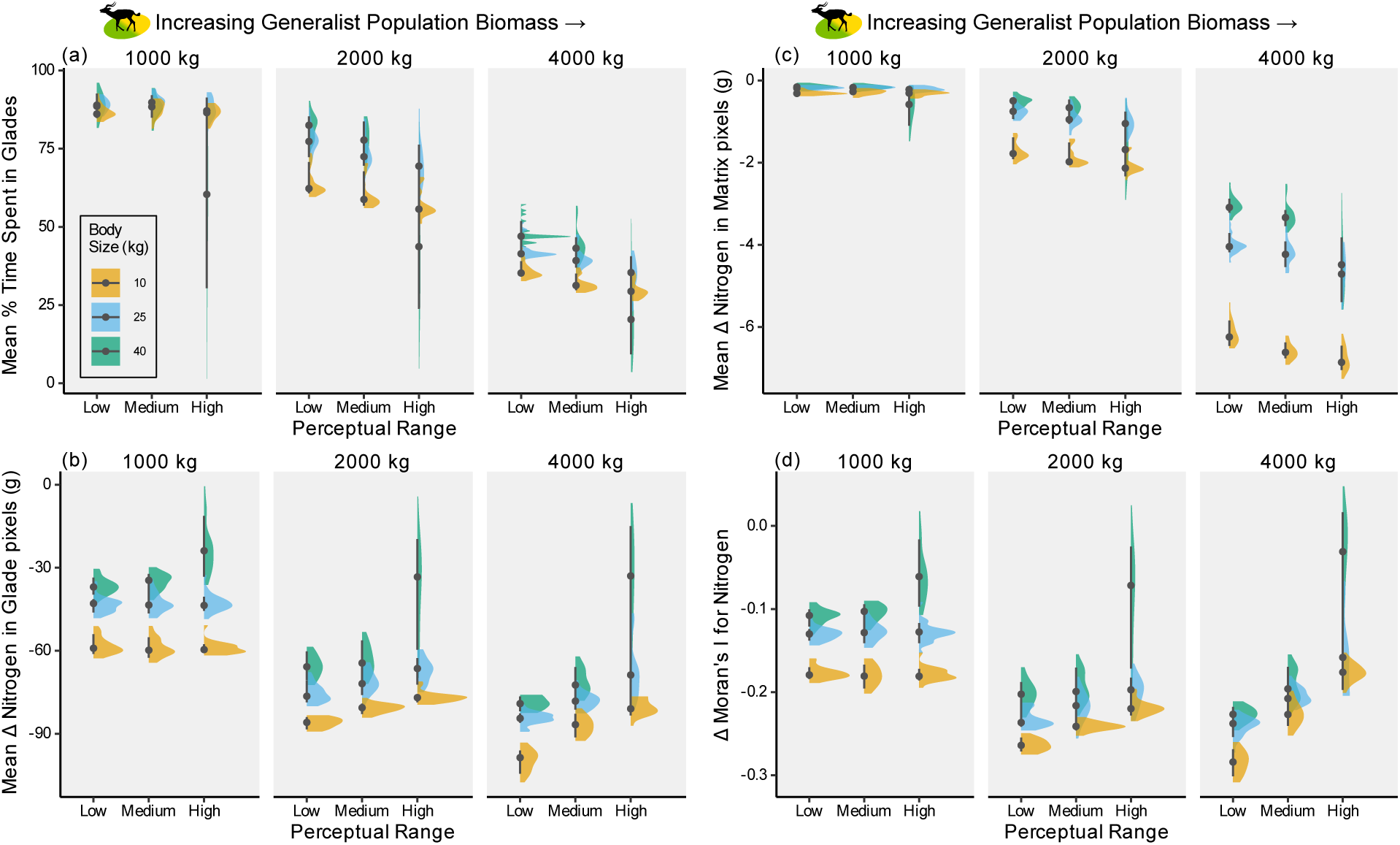
From scenarios placing generalists of varying body sizes, population biomass, and perceptual ranges on the landscape, distributions of data for resulting (a) time that the generalists spent in Glades, (b) net Nitrogen loss from Glades, (c) net Nitrogen loss from Matrix, and (d) change in Moran’s I for Nitrogen across the landscape. Black points show median across 25 replicate simulations per parameter combination for three different parameter combinations, and black lines show distribution of 75% of the data. The “Medium” perceptual range corresponds to the parameterization shown in Figures 2-5; the “Low” category represents a parameterization with half of the “Medium” perceptual range, and the “High” category represents twice the “Medium” perceptual range.

In the previous sections, we showed that at low population biomasses, generalists did not sufficiently exploit Glades to cause them to leave and spend significant time in the Matrix. Here, we found that increasing generalists’ perceptual ranges did not substantially alter their (minimal) impacts on Matrix resources when population biomass was low (**Fig. 6b**). In these scenarios, only at the largest body size did increasing perceptual range shift generalists’ impacts on N spatial correlation (**Fig. 6d**). Thus, we found that an interaction with population biomass determined the impact of perceptual range on spatial resource distribution.

### Sensitivity testing

The patterns of spatial change in N caused by consumers of different body sizes and population biomass remained qualitatively the same when the neighborhood size over which Moran’s I was calculated increased from only the nearest neighbor to the 4, 10, and 17 nearest neighbors, although reductions (by generalists) and increases (by specialists) in spatial autocorrelation were less accentuated as neighborhood size increased (**Fig. S4**). Additional simulation scenarios showed that increasing the proportion of consumed nitrogen that was deposited by consumers from 50% to 75% also did not substantially change these results (**Fig. S5**). In further sensitivity testing, we found that unlike varying the perceptual range, neither increasing the frequency of nutrient deposition by consumers (**Fig. S7**) nor altering the mortality rate (**Fig. S8**) resulted in different spatial patterns than our primary parameter set.

## Discussion

The role of animals in structuring resource distributions in space is poorly understood. Here, using a spatially-explicit, individual-based model, we show that organismal traits can make the difference in whether consumers amplify or erode spatial resource heterogeneity. We found that a key trait determining whether consumers amplify or erode resource patchiness is species’ preference for the different habitats, and that larger perceptual ranges reduced the extent to which generalists eroded landscape structure. Patchy landscapes have driven the development of ecological theory (Macarthur and Wilson 1967, Hanski 1999, Loreau et al. 2003a), yet to date, models assessing the effects of animal movement, foraging, and resource deposition have primarily focused on landscapes without spatial structure in resource availability (Earl and Zollner 2014, Bampoh et al. 2019, 2021, Ferraro et al. 2022). When spatial structure is included, over seasonal time scales, we showed that specialists on lower-quality (Matrix) habitat amplified existing resource heterogeneity between the resource hotspots (which they avoided) and the low-quality habitat. Conversely, generalists who could take advantage of high-resource Glades eroded the landscape’s spatial structure.

### Potential homogenization of resource landscapes

The seemingly obvious outcome that specialists increase and generalists erode spatial resource heterogeneity nevertheless provides an important insight in an era of global change because generalists often thrive where specialist species are more vulnerable to habitat loss and fragmentation (Clavel et al. 2011, Keinath et al. 2017). Additionally, the Anthropocene is characterized by a proliferation of non-native species, and generalism is associated with invasion success (Romanuk et al. 2009, Ord and Hundt 2020). Our results suggest that replacement of specialists with generalists could lead to more spatially homogenous resource distributions across landscapes over short timescales. This homogenization would likely reduce both spatial clustering of nutrients, as well as dissimilarity of the plants that serve as food resources. Evidence from temperate forests already shows that abundant generalist herbivores can reduce beta diversity of plant communities (Holmes and Webster 2011). The potential for landscape homogenization that we show here could further disadvantage specialist species and reduce landscape resilience to environmental impacts.

Larger body sizes, and associated larger perceptual and movement ranges, may mitigate this trend towards homogenization to some extent. We found that consumers with greater movement ability could travel with larger step lengths, and both their consumptive and depositional effects were therefore less spatially autocorrelated. We also found that as generalist consumers’ sensing and movement ability increased relative to their body size, they spent more time traversing the landscape and less time in the Glades than when their sensing and movement ability was lower. In our model, the distance over which consumers can sense the landscape in a timestep and the distance they can move over in that same timestep, are controlled by the same parameter; this assumption is not necessarily true for all species, and neglects the role of social information in finding the best foraging movements (Little et al. 2022). But here, the net outcome of the sensing and movement processes was that with relatively larger perceptual ranges and movement abilities, generalists did not as significantly erode the nutrient heterogeneity set by the pre-existing landscape structure over the course of a single growing season.

Individuals with the largest perceptual ranges can glean more information: they are closer to “perfect information use”, which changes how quickly and efficiently they can find and move to areas of the landscape with the best resources, consume those resources, and contribute to ecosystem processes (Little et al. 2022). Perhaps counterintuitively, this can lead to more possible behaviors amongst individuals compared to those constrained by small perceptual ranges. With limited information use, these latter individuals are more likely to spend time near local optima. By contrast, in our model, large consumers with large perceptual ranges could often sense multiple high-resource Glades to choose amongst. In nature, this could lead to several different strategies. Individuals could, for example, stay in one resource-rich spot where forage is good, avoiding the energy cost of movement, or they could move frequently, leaving the current location as soon as the glade next door seems better. The optimal choice may depend on what other information the individual has access to: their condition, the behavior and location of conspecifics, the location and likelihood of other risks, and more. This choice of strategies would not be available to individuals with weaker perception, whose only options may be one resource-richer area or the matrix.

In our model, different behavior by individuals with large perceptual ranges may also have emerged from the different random configurations of Glades when the landscapes were initialized. If several Glades are nearby but one is farther away, the population may predominantly forage in the region with more Glades, which according to the marginal value theorem would be substantially better than the rest of the landscape (which has proportionally more lower-quality Matrix). While this would homogenize the region, it could protect landscape-scale heterogeneity due to less grazing in the far-away Glade (and its surrounding Matrix). Meanwhile, if Glades are more uniformly distributed across the landscape, the landscape regions would be similar according to the marginal value theorem. In this case, consumers in all parts of the landscape would likely switch between Glades as forage is consumed, leading to consistent reductions in N across Glades and an overall more homogenous resource landscape at the end of the season. These two differing outcomes based on initial landscape configuration could contribute to the highly variable degree of homogenization that we observed when large-bodied consumers have high perceptual ranges (**Fig. 6d**).

Our results also suggested a non-linear effect of generalists’ perceptual range on resource spatial distributions. For larger perceptual ranges to serve as protection against the erosion of spatial heterogeneity, we found that either perceptual ranges must be very high (for example, with both the highest perceptual range scaling parameter and the largest consumer body size), or the consumer population biomass must be high enough for the more moderate effects of a smaller increase in perceptual range to accumulate through the activity of many individuals in the population. Perceptual range and movement distances are shrinking due to global change, both by constraining species movement and habitat use via barriers and habitat conversion (Tucker et al. 2018), and due to the aforementioned reduction in body size, and the link between body size and foraging scales (Straus et al. 2024). The concurrent shifts towards generalism, smaller body sizes, and associated smaller perceptual ranges could result in drastic erosions in spatial heterogeneity. Our results suggest an additional motivation for protecting large consumers and their ability to range over great areas of their landscapes.

### Importance of species traits in shaping resource heterogeneity

The specialists in our simulations provided an example of how risk avoidance strategies (here, avoiding resource-rich Glades) can reinforce spatial structure and patchiness of landscapes by strengthening existing resource gradients (for an empirical example, see Germain et al. 2013). In addition to accentuating the difference between Glades and Matrix habitat, these same specialists also created finer-scale spatial resource heterogeneity where it was previously lacking. Our landscapes featured no initial spatial clustering of resources *within* a habitat type, but over the course of our simulated growing season, spatial resource heterogeneity in the Matrix increased. This was because the removal of N by foraging followed individuals’ movement patterns, which were spatially autocorrelated, particularly when consumers are small and can both perceive and move only over short distances. This increase in fine-scale heterogeneity was not captured by Moran’s I, which better reflected the larger differences in N concentrations between Matrix and Glades, but is visible as waves of resource concentrations in landscape depictions (**Fig. 1**, “150 Large Specialists”). Thus, the effect of organismal activity on landscape-scale nutrient spatial heterogeneity depends on both the existing structure of the landscape (that is, the initial spatial distribution of resources) and the scale of observation (that is, the spatial grain at which one is measuring heterogeneity – for example, the habitat or the landscape).

Globally, many species of large-bodied vertebrates are experiencing range contractions, whereas smaller-bodied vertebrates are more likely to be experiencing range expansions (Pacifici et al. 2020). The largest consumer species are also at higher risk of extinction than are their smaller counterparts (Enquist et al. 2020). This raises the question of how a reduction in body size will affect nutrient cycling and distribution on landscapes. For the same total biomass, smaller consumers consume more forage due to higher mass-specific metabolic needs (Brown et al. 2004). Our results imply that in patchy landscapes, the impact of species turnover from large to small generalists arises not only from this metabolic scaling, but also depends on both initial community biomass (i.e., the summed biomass of all individuals in the population) and whether shifts in body size are accompanied by an increase or decrease in total biomass. At small body sizes, larger populations of generalists homogenized the landscape more than smaller populations by depleting resource hotspots. This is somewhat in contrast to previous findings that high densities of consumers can create spatial structure when competition leads to overexploitation of resources (cold spots) and carcass deposition leaves resource hotspots. Mortality may not have been high enough in our model to create many hotspots, and this creation of heterogeneity may be more apparent at larger body sizes, for example with caribou (Ferraro et al. 2022). We speculate that with small consumers – especially those even smaller than the ones modeled in our study, such as rodents – hotspots may be smaller (due to smaller carcasses) and more evenly distributed on the landscape when there is a high density of consumers with small movement ranges. This could limit the spatial structuring of resources by small consumers.

At larger body sizes and greater total biomass, the further depletion of resource-rich Glades by a larger population was offset by the increased consumption of resources in the lower quality habitat and spatial homogenization plateaued, and in some cases spatial structure even increased again as consumers created heterogeneity in the Matrix due to their spatially autocorrelated foraging. This provides further evidence that conservation of large-ranging species should be an urgent priority. It is important to note that even if declines in landscape-scale resource clustering halt when some threshold is reached (when initially higher- and lower-resource areas become somewhat equal), increases in population size or total biomass, and shifts from large to small body sizes, would increase overall net rates of consumption and recycling of resources, and could have profound consequences for ecosystems.

### Limitations and considerations for future work

Our model did not include population dynamics of either consumers (beyond mortality) or forage resources, and only considered the spatial dynamics of net resource availability. Our model also ran over the course of a single growing season, exploring the ways that consumers could alter the environmental template; over larger time scales, legacy effects, plant population dynamics, and the complexities of plant-soil-consumer nutrient cycling could either dampen or intensify the short-term restructuring we examined here. However, the simplification and short time scale that we implemented in our modeling approach increased computational feasibility and enabled us to isolate the impact of vertebrates on nutrient distributions. Our results show that these direct effects can be substantial, even over short timeframes. However, feedbacks with population dynamics should be explored, and could contribute to determining how long-lasting the effects on nutrient distributions might be. Previous work using non-spatially-explicit approaches has shown that seasonal resource heterogeneity stabilizes consumer populations, if the lower-quality resource (habitat, in our conceptualization) is nonetheless palatable and can be used as a “buffer” resource (Owen-Smith 2004), similar to how the generalists choose resources in our models.

Another caveat is that in our model, not all N consumed in forage is returned to the landscape. Some N is incorporated into consumer bodies, and consumer death was not high enough in our model to offset this; even doubling the mortality rate did not do so, and only minimally affected the overall results. This dynamic is not unrealistic over the course of a single wet season, based on mortality estimates from the literature (Ogutu et al. 2012, Owen-Smith and Goodall 2014), however models that run over longer timescales, including the dry season when conditions are more harsh, may show a greater impact of mortality. We also did not explicitly model processes such as predator and scavenger export of nutrients (which would likely occur over much larger spatial scales than our entire simulated landscape; white-backed vultures, for example, can forage over hundreds of kilometers (Phipps et al. 2013)), leaching, or nutrient volatilization, instead incorporating assumptions about how much N was returned to the landscape by excretion, egestion, and carcass deposition.

## Conclusions

Here, we showed through simulations that animals have the ability to shape resource heterogeneity on the landscape over short periods of time such as a single season, thus shifting the environmental template for subsequent consumer-resource interactions. Landscape heterogeneity is important because it creates different niches that can serve to sustain biodiversity (Ryser et al. 2021). Our results suggest that species’ traits and abundance are key mediators of their effect on resource distributions, which could lead to varying impacts of future biodiversity change. Despite simplifications, our finding that generalists (which we based on impalas) homogenize N distributions by reducing N in Glades aligns with empirical results in an intriguing way: impalas increase the relative density of a well-defended plant species (Ford et al. 2014) which has less foliar N than a poorly-defended congener (Coetsee et al. 2023) that is abundant where impala are rare, thus altering spatial heterogeneity of N in plants. This more mechanistic example shows how consumers can shape the functional distribution of resources across landscapes. We see great potential for further integrating modeling with empirical findings to improve our understanding of animals’ influence on nutrient spatial dynamics.

We propose two future directions for spatially-explicit, agent-based modeling to explore the impacts of these projected changes. The first is to explicitly model different chemical forms of nutrients, which are more or less bioavailable to organisms (Hempson et al. 2015). This approach would account for the fact that nutrients deposited through dung and urine are not immediately available to consumers, but must first be transformed into plant shoots. We expect that coupling these considerations with longer simulations that account for consumer population dynamics would lead to new spatial patterns of foraging movement by consumers, and thus new emergent dynamics of nutrient spatial distributions. Second, further work should also explore how competing consumers and trophic interactions modify the patterns found in this study. For example, we modeled species based on known risk-avoidance strategies rather than including predators in the models, but doing so may lead to behavioral shifts and emergent dynamics, additionally because trophic role influences resource deposition (Mackay et al. 2021). Furthermore, consumers exist in multi-species communities and trait-based differences among co-occurring species could lead to complementarity of resource use, altering the patterns that we found here. Both of these future directions could be explored using starting landscapes that are both spatially homogenous and those with spatial structure, including not only among-habitat spatial structure (which we have modeled here with Glades and Matrix), but also within-habitat spatial structure, which is also important to ecosystems and consumers. Incorporating additional ecological complexity into spatially-explicit models could enable a better understanding of how organisms will shape their own fates via resource dynamics in future landscapes.

## Data / code availability

Data and code are available to peer reviewers at https://zenodo.org/records/14290828.

## Acknowledgments

This research was enabled in part by support from the BC DRI Group, Simon Fraser University’s Supercomputer Cedar (www.sfu.ca/research/supercomputer-cedar), and the Digital Research Alliance of Canada (www.alliancecan.ca). We thank Matteo Rizzuto and members of the Little Ecology Group for valuable feedback on the manuscript. We appreciate the time spent by three anonymous reviewers, whose comments greatly improved this paper.

