## Appendix I for "Spatial signature of resource distribution is mediated by consumer body size and habitat preference"

**Contents:**

**Appendix I - Model Overview, Design, and Details** 2

Includes Figure A1, Tables A1 & A2

**Appendix II - Supplementary Figures & Tables** 12

Includes Table S1, Figures S1-S8

**References** 23

### Appendix I - Model Overview, Design, and Details

This appendix presents a model description using the Overview, Design, and Details framework (Grimm *et al.* 2010).

#### Overview

*Purpose*

The purpose of this model is to examine how the dynamics of nutrients (specifically, nitrogen) are determined by animals in a spatially-explicit landscape made up of a mosaic of habitat types with different resource availability. Agents represent animals that select landscape areas in which to consume resources. Agents navigate this landscape based on the nutrient content available on different pixels. They also excrete/egest nutrients as they move, and if they die the nutrients contained in their carcasses are also deposited on the landscape. Nitrogen loss (through consumption) and gain (through deposition) are tracked through time in each grid cell in the landscape. The model is used to ask the following questions: how do

1) Animal body size;
2) Number of animals;
3) Habitat preference; and
4) Aggregation type (i.e. herding vs. solitary lifestyles)

affect patterns of net nitrogen gain and loss across the landscape?

*Entities, state variables, and scales*

The model is built in NetLogo (Wilensky 1999) and has two types of entities: grid cells in the landscape ("pixels") and agents. The landscape consists of a 141x141 grid of pixels, each of which represents a 10x10m area of a terrestrial landscape. The 1.9881 $km^2$ extent of our landscape is slightly larger than the upper estimates of home range size for herds of female impalas (Murray 1982), the larger of the two herbivores that inspired this study.

Pixels are wrapped on a torus so that agents can move around the landscape without creating edge effects. An alternative way of avoiding edge effects would be to implement a reflective boundary so that rather than passing to the other side of the landscape, consumers move back in the direction from which they arrived when they reach the boundary. Both approaches involve assumptions about the landscape. A reflective boundary assumes that the organisms are truly limited by the spatial extent of the modeled landscape, and can create some edge effects by increasing time spent in the pixels directly surrounding the boundary. Meanwhile, the torus simulates a scenario where as an organism leaves the landscape, another enters, and it spreads the occupancy of pixels near the boundary across each side of the boundary rather than reflecting the consumer back in the direction it came from. Importantly, both approaches are the same in terms of preserving total population size. Because we were not modeling a particular exact landscape, and because the home range of organisms typically varies with body size and we systematically tested different body sizes in this model, we chose to use a torus.

There are six total classes of agents: specialist animal, of generalists animals, and feces and carcasses for each of two species. Animal agents move across the landscape and create feces and carcasses which remain in the pixel where they were created. Each time-step of the model represents 3 hours of a day, with the behavior of animal agents varying depending on time of day based on empirical daily activity budgets. The model is run for 720 time steps or 90 days, the rough length of a season.

*Pixels.* Pixels are characterized as one of two habitat types, which are loosely based on Kenyan landscapes featuring "Glades" (more productive glades that are former cattle corrals) and "Matrix" (Augustine 2003, 2004; Veldhuis *et al.* 2018). Glades are circular areas with higher biomass of forage, and more nitrogen-rich (e.g. nutritious) forage in a Matrix of lower-forage, lower-nitrogen grassland. For each pixel, forage availability, the number of times a pixel has been grazed, the nitrogen removed by that grazing, the number of times agents have deposited feces/urine in the pixel, and the nitrogen deposited through feces/urine are tracked.

*Agents*. The model focuses on behavior of agents, particularly of specialists and generalists. Specialists prefer the Matrix grassland habitat and do not enter the Glades; this species approximates Guenther's dik-diks, which are pure browsers, do not eat grass, and prefer bushy habitat for predator avoidance (Augustine & McNaughton 2004; Kingswood & Kumamoto 1996). Grazers are a more generalist species which can use either habitat; this species approximates impala, which avoid the bushier matrix habitat in a completely different predator avoidance strategy and both browse and graze the more nutritious Glades (Augustine & McNaughton 2004; Ford *et al.* 2014). Energy requirements and energetic cost of movement scale allometrically in the same way for agents of both species, however their movement is different (see *Navigate* submodel). Agents of each species move across the landscape, consuming forage (and the nitrogen contained therein) in pixels at a rate that depends on their energetic needs and the amount of available forage. The agents of these two species accumulate nitrogen in their bodies based on feeding, and deposit feces containing a percentage of that nitrogen back onto pixels (see *Poop* submodel). When they die, either stochastically or of hunger, they deposit carcasses (and the nitrogen contained therein) on pixels (see *Nutrient transfer to pixels* submodel).

Feces are hatched by specialist or generalist agents, see (see *Poop* submodel), and have characteristics of the amount of nitrogen contained therein, as well as the timestep in which they were created. Here, we refer to these as "feces" for simplicity, but the deposits are intended to represent the combined deposition of both feces and urine (that is, both excretion and egestion). Carcasses are likewise hatched by specialist or generalist agents, see (see *Navigate* submodel), in this case when they die, and have characteristics of the amount of nitrogen contained therein, as well as the timestep in which they were created.

*Process overview and scheduling*

The model is initialized and then run for 720 timesteps (**Figure S1**).

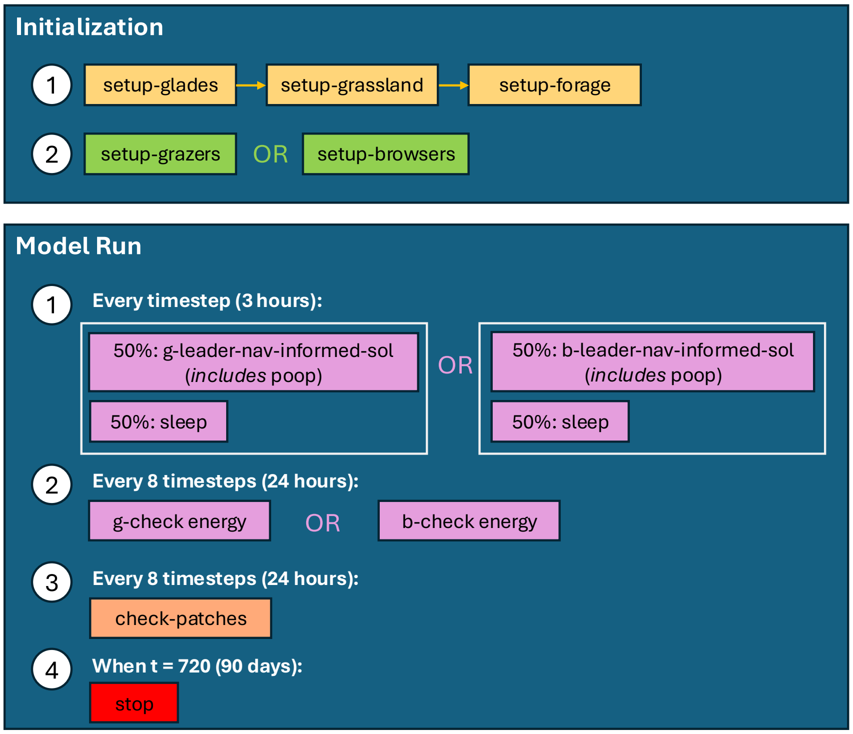


**Figure A1**. The order of submodels during initialization and setup, when run for only one species on the landscape and in the solitary behavioral mode. Glades are placed on the landscape first ("setup-glades"), then Matrix ("setup-grassland"), and then the amount of forage is chosen for each pixel based on its habitat type ("setup-forage"). Based on user-defined parameter choices, either generalists ("setup-grazers") or specialists ("setup-browsers") are placed on the landscape. During the model's running, some submodels are run every timestep while others are run less frequently. Every timestep, each consumer either runs through the "leader-nav-informed-sol" or the "sleep" module; the navigation modules are slightly different for generalists (signified by g) and specialists (signified by b). The navigation module includes the poop module. Every 8 timesteps, the "check energy" module is run, which controls mortality events, and the "check-patches" module disperses N from carcasses deposited during the previous 24 hours onto the landscape. When the model reaches 720 timesteps, it stops.

Initialization of the model begins by choosing which species is present on the landscape, and that species' aggregation type, total biomass, and average individual body mass. This parameter initialization can be done either through the NetLogo GUI, or through an R interface; we used the 'nlrx' package (Salecker *et al.* 2019). The landscape is then built with a random configuration of Glades, and agents of the species are randomly placed on the landscape. Specialists are placed in Matrix pixels, and generalists are placed in Glades. If the aggregation type is set to individual, then agents are each randomly placed; if the aggregation type is set to herd, then a first individual of the species is randomly placed, and then the others are randomly placed within a radius of that leader to form a herd.

For each timestep,
(1) generalist or specialist agents check whether there is a leader - if in a herd scenario and the leader has died, a new leader is randomly assigned, while in any scenario if there are no living agents remaining, then the simulation ends;

(2) the mean nutrient content of the landscape as a whole and of the matrix are calculated;
(3) generalist or specialists agents undergo either the *Sleep* submodel or the *Navigate* submodel, each with a 50% probability, with the *Navigate* submodel including the *Poop* submodel;
(4) one unit is added to the timestep counter.

At the end of each day (every 8 timesteps),
(1) the *Mortality* submodel is performed;
(2) the *Nutrient transfer to pixels* submodel transfers the nitrogen contained in carcasses deposited during those 8 timesteps to the pixels where they are located;
(3) one unit is added to the days counter.

At the end of the growing season (90 days), the simulation is halted.

### <u>Design Concepts</u>

*Basic principles*

Animals consume resources in order to survive. Their movement and activity on landscapes is both a consequence of resource heterogeneity - that is, they can consume resources where they are present, but not where they are low or absent - and a cause of it, as they redistribute resources through egestion, excretion, and carcass deposition. The model was designed to investigate how animals of different body sizes, sociality types, and abundances affect resource heterogeneity at the landscape scale through their sensing of resource distributions, their movement, and their activity.

*Emergence*

Whether a pixel is grazed or not, how many times, and the net nitrogen gain or loss are results emerging from the behavior of the animal agents. In particular, spatial patterns in these pixel-level variables are the outcome of interest.

*Adaptation*

Animal agents have a combination of fixed and adaptive behaviors. As an example of a fixed behavior, submodels for eating and resting are run probabilistically depending on the proportions of hours in a day typically spent eating and resting by impala and dik-dik (see *Sleep* submodel), and even if an agent has not consumed enough energy to survive they cannot make up for this by eating in a time-step that is designated for rest. Similarly, the amount of forage in a pixel that is available to them to eat is capped at 50% of the forage present in a pixel, regardless of the agent's hunger level. On the other hand, an example of adaptive behavior is that animal agents' movements are based on sensing the environment in a way that is designed to maintain or improve fitness (see Objectives, below). Foraging pixels are chosen by balancing energy costs (the distance that would have to be traversed to reach them) and energy gain (the amount of forage available in the pixel): when there are pixels within an agent's perceptual range with better-than-average forage, they move to the closest one.

*Objectives*

For animal agents, the objective is to consume sufficient forage to survive and to accumulate the most nitrogen while doing so. The metabolic needs of each agent is calculated based on its body mass, based on the following equations: daily needs in kJ = 7.94 * (body mass in g) ^ 0.646, and 11.5 kJ of energy per gram of dry matter (Nagy 2001).

*Learning and prediction*

Objectives and behaviors used to meet them remain the same during the simulations.

*Sensing*

Specialist and generalist agents sense nutrient levels in the pixel where they are currently located; sense nutrient levels in pixels in a set radius from their location, with radius based on body size (range (in meters) ~ 5 * bodymass (in kg)); and sense the mean nutrient levels of the landscape. They also sense whether there is another individual on a given pixel in that same set radius. In herd scenarios, "followers" sense where the leader is located.

*Interaction*

Specialist and generalist agents interact with each other in the model by avoiding pixels where another agent is already located. In herd scenarios, there is an additional layer of interaction where agents follow a leader. Specialist and generalist agents interact with pixels by consuming forage from them, and depositing feces and carcasses onto them.

*Stochasticity*

There are several stochastic elements of the model. At the beginning of each simulation, pixel forage amounts and specialist or generalist agents' body sizes are each drawn from normal distributions around a set mean and variance. The location of Glades in the pixel landscape and the initial placements of agents on the pixels is random. Where forage is good, specialists and generalist agents move to the closest good pixels, but when they are in an area with insufficient forage or worse-than-average forage, stochastic elements are introduced to their movement in order to help them reach new areas of the landscape outside their perceptual range (see *Navigate* submodel). Feces are hatched semi-stochastically, with a probability of 33% if the agent has not defecated in more than two timesteps (6 hours), however feces are hatched with 100% probability if the agent has not defecated in more than six timesteps (18 hours). Mortality occurs with a fixed probability per population size (see *Mortality* submodel), but the individual agent of a species selected for stochastic mortality is chosen randomly.

*Collectives*

In some runs of the model, the specialist or generalist agents can form a herd, which move across the landscape within a certain radius of one another. This behavior emerges based on individual behavior, and the herd has no behavior or state variables of its own. Instead, the leader makes decisions about where to forage, and the followers move based on the behavior of the leader.

*Observation*

At the end of each 90-day run, all data from all pixels is recorded. Spatial and non-spatial statistics on these data can then be calculated based on the user's research questions.

### **Details**

*Initialization and Input data*

The user designs a modeling scenario by choosing values of the parameters described in Table A1:

**Table A1.** User-defined inputs.

| Variable | Possible values | Explanation |
|---|---|---|
| Grazer-aggr-type | Herd or solitary | Whether generalist agents live/move in herds, or whether individual agents independently devise their own movement to meet objectives. |
| Total-grazer-biomass | Positive integer | The total amount of generalist agent biomass placed on the landscape at the beginning of the simulation, in kg. |
| grazermass | Positive integer | Mean body mass of generalist agents (kg) |
| browser-aggr-type | Herd or solitary | Whether specialist agents live/move in herds, or whether individual agents independently devise their own movement to meet objectives. |
| Total-browser-biomass | Positive integer | The total amount of specialist agent biomass placed on the landscape at the beginning of the simulation, in kg. |
| browsermass | Positive integer | Mean body mass of specialist agents (kg) |

If Total-grazer-biomass is set to 0, then no generalist agents are placed on the landscape. Likewise if Total-browser-biomass is set to 0, then no specialist agents are placed on the landscape.

The landscape is initialized by setting pixels to be either Glades or grassland Matrix. To set up circular Glades, 4 random pixels are chosen to be glade centers. 5% of the landscape is targeted to become Glades, with an average Glade radius of 90 meters, slightly larger than empirical estimates of bomas' average size by Augustine (2003). To achieve this, every pixel in a radius of 9 from one of these Glade centers is then also made into a Glade pixel. The landscape is then checked to see whether the overall proportion of Glade pixels reaches 5%; if two Glade centers are placed within 9 pixels of each other, for instance, they will form one single multicenter Glade with less area than that of two individual Glades. If needed, more Glades are added by randomly selecting additional Glade centers, and then the landscape is checked again until the threshold proportion of Glade pixels is reached. The amount of forage for each pixel is then generated by drawing randomly from a normal distribution centered on 50 g/m$^2$ for Glades and 33 g/m$^2$ for Matrix, with forage containing 6% nitrogen (N) in Glades and 4% in Matrix.

The number of agents placed on the landscape is determined by dividing the user-defined total desired biomass of generalists or specialists by the user-defined mean body size of that type of agent (see **Table A2**). Specialists are placed in the grassland Matrix, and generalists are placed in the Glades. In simulations where agents of a given species act individually ("solitary"), they are all set to be leaders and placed randomly on the landscape within their designated habitat type. In simulations where agents of a given species are in a herd, one agent is randomly placed on the landscape within its designated habitat type and set as a leader. Then, the subsequent agents are placed on the landscape within their designated habitat type, in a random location that is within the "herd-radius" distance from the leader, calculated based on the number of agents of that species (see **Table A2**). These agents are designated as followers, and the leader is designated as their leader. Perceptual range is defined as a radius that scales linearly with body size: the radius in which an agent can sense is, in pixels, 0.5 times the average body size (in kg) of specialist or generalist agents, and is the same for all agents of a given species regardless of their realized body size (see **Table A2** for how this produces an area-based perceptual range, which scales quadratically with body size).

**Table A2** Selected global variables.

| Variable | Value | Explanation |
|---|---|---|
| n-grazers-to-create | total-grazer-biomass / grazermass | The number of generalists agents to place on the landscape at initialization |
| n-browsers-to-create | total-browser-biomass / browsermass) | The number of specialist agents to place on the landscape at initialization |
| Grazer-herd-radius | √ (3 / π) * n-grazers-to-create | For generalists, when grazer-aggr-type is set to "herd", the radius of the herd size (that is, within what distance "follower" agents should be to the "leader" agent); set so that for every agent in the herd, there are 3 pixels within the herd's occupied area (their current pixel and two empty pixels) |
| Grazer-omniscience | Maximum of either 2 or 0.5*grazermass | Perceptual range: 10's of meters over which generalists can select habitat; set at 5 meters (0.5 pixels) per kg of body mass, or 2 pixels away, whichever is larger. As a result, the area of a generalist's perceptual range (in km$^2$) scales quadratically with consumer body mass, according to the equation<br>Perceptual range = $\pi$ * (0.5 * body mass)$^2$ *100 |
| Browser-herd-radius | √ (3 / π) * n-browsers-to-create | For specialists, when browser-aggr-type is set to "herd", the radius of the herd size (that is, within what distance "follower" agents should be to the "leader" agent); set so that |

| | | |
|---|---|---|
| | | for every agent in the herd, there are 3 pixels (their current pixel and two empty pixels) |
| Browser-omniscience | Maximum of either 2 or 0.5*browsermass | 10's of meters over which specialists can select habitat; set at 5 meters (0.5 pixels) per kg of body mass, or 2 pixels away, whichever is larger. As a result, the area of a specialist's perceptual range (in km$^2$) scales quadratically with consumer body mass, according to the equation<br>Perceptual range = $\pi$ * (0.5 * body mass)$^2$ *100 |
| mean-nutrients-matrix | [Calculated at the end of each day] | The mean nutrient content of all Matrix pixels in the landscape |
| mean-nutrients-glade | [Calculated at the end of each day] | The mean nutrient content of all Glade pixels in the landscape |
| mean-nutrients-overall | [Calculated at the end of each day] | The mean nutrient content of all pixels in the landscape regardless of habitat/pixel type |

*Input*

This model does not incorporate input data.

*Submodels*

<u>Sleep</u>

In each timestep, specialist and generalist agents have a 50% chance of resting; this is based on empirical findings that both dik-diks and impala spend roughly 50% of their time feeding (Manser & Brotherton 1995; Owen-Smith & Goodall 2014). In this case, they remain in the pixel where they are currently located and nothing further is calculated or acted upon, other than increasing that agent's time-since-pooped counter (see *Poop* submodel).

<u>Navigate</u>

Navigation is the most complex submodule and the one from which most landscape patterns emerge. If navigating, specialist and generalist agents must find a new pixel; agents do not stay in the pixels where they are located and continue to eat there. The navigate submodel works differently for specialists, which only navigate in the Matrix pixels, and for generalists, which navigate across the whole landscape. For each species, there are also three different models depending on their identity: individuals, leaders of herds, and followers of herds. These sub-submodels are b-leader-nav-informed-sol (for solitary specialists), b-leader-nav-informed-herd (for specialist herd leaders), b-follower-nav-informed (for followers in specialist herds), g-leader-nav-informed-sol (for solitary generalists), g-leader-nav-informed-herd (for generalist herd leaders), and g-follower-nav-informed (for followers in generalist herds). In all cases, the agents operate roughly according to the marginal value theorem, which states that a consumer should leave its current location if the resource level falls below the marginal resource level for that habitat type (Charnov 1976). Thus, agents compare the N available to them in forage to the mean N in forage at the level of the entire landscape (for generalists) or entire matrix (for specialists), and then either select a nearby pixel to move to if the marginal value is high, or if the marginal value is low they make a larger movement in order to access a new part of the landscape which may have better resources. Thus, the ways in which agents navigate depend on the species and the agent's leader/follower identity as follows:

*Solitary agents:* Check if the average nutrient content of pixels in their perceptual range is better than the landscape average. If so, they move to the closest pixel in their perceptual range which contains better-than-landscape-average nutrients, and no other generalist or specialist agent present; if all better-than-average pixels have other agents present, then they simply move to the closest open pixel. If the pixels in their perceptual range are not better than the landscape average, then the agent moves to the closest pixel outside their perceptual range that does not have another generalist or specialist present. This type of movement is meant to represent longer-distance, exploratory journeys into unknown habitat with the potential to reach areas of higher resource availability. Note that for specialists, the procedure just described is only applied to Matrix pixels, but for generalists it considers all pixels.

*Herd leader:* The navigation proceeds as for solitary agents, except that the range of area considered expands to encompass the perceptual range of all agents in the herd (i.e., herd-radius + perceptual range). This is meant to mimic social information use within a herd, without adding the model complexity and computing time to mechanistically model information transfer. Once again, for specialists, the procedure just described is only applied to Matrix pixels, but for generalists it considers all pixels.

*Herd followers:* Herd followers check whether there are any pixels within the herd-radius from their leader which contains better-than-landscape-average nutrients, and no other generalist or specialist agent present. If so, they move to the closest one. If not, they simply move to the closest pixel in the herd-radius that has no other specialists or generalists present. If there are no pixels within the herd-radius without another agent present (note: this should only occur for specialists, in the case that part of the area in the herd-radius is Glade which they will not enter), then the agent moves to the closest open pixel, even if it is outside their herd-radius. Once again, for specialists, the procedure just described is only applied to Matrix pixels, but for generalists it considers all pixels.

In all movement cases, the distance traveled to get to this pixel is added to the distance-traveled counter for the agent. The agent then checks if their new pixel has enough forage available to meet their energy needs (which are calculated as 25% of their daily-forage-need, since there are 8 timesteps per day and a 50% chance of feeding at each timestep). If so, they consume that amount of forage. If not, then they consume 50% of the forage present on the pixel. This reflects that not all biomass in a given location is palatable/available to all consumers (that is, herbivores cannot graze a pixel to completely bare ground), and also that there is competition with species not explicitly modeled in our scenarios (Buchmann *et al.* 2011). In either case, the amount of forage actually consumed is added to the net-forage-consumed-grassland or net-forage-consumed-matrix counter for the agent; the nitrogen consumed therein is added to the nitrogen-to-poop counter. For the pixel, the forage consumed is subtracted from the forage-mass-current, the time-since-eaten is set to zero, and the times-grazed is increased by one. N-current is recalculated to subtract the amount of nitrogen present in the forage that was consumed. At the end of the foraging process, the *poop* submodel is called (see next section).

Poop

For both specialist and generalist agents, the *poop* submodel is run as part of the *navigate* submodel (that is, it does not occur for agents that are resting in the *sleep* submodel). This submodel first asks, for an individual agent, whether it has been more than two timesteps (six hours) since the agent last excreted/egested. If so, then the agent may hatch a new poop agent. If it has been between two and six timesteps since the last poop agent was created, then there is a 33% chance of the specialist or generalist agent going through this process to create a new poop agent. If it has been greater than six timesteps (18 hours), then there is a 100% chance that the agent hatches a poop agent. Whenever this occurs, the pixel-level variables times-pooped, net-N-deposited, and either net-N-pooped-glade or net-N-pooped-grassland (depending on where the agent is located) are updated, either by adding one to the times-pooped counter or by adding the nitrogen contained in the new poop agent to the net-N counters. Then, the time-since-pooped and nitrogen-to-poop are set to 0 for the specialist or generalist agent. If the agent did not hatch a new poop agent in this timestep (for example, if it has been 2 or fewer timesteps since the last poop event, or if it has been 3-6 timesteps but this was one of the 66% of times that they did not poop), then the time-since-pooped counter is increased by one.

Poop agents contain 50% of the nitrogen consumed in forage since the last time the specialist or generalist agent pooped; this is based on empirical data of N deposited / N consumed by ungulates in Yellowstone National Park (Singer & Schoenecker 2003). The poop agent is placed on the pixel where the specialist or generalist agent is currently located.

Mortality

The mortality submodel operates slightly differently for specialists (the b-check-energy sub-submodel) and generalists (the g-check-energy sub-submodel). Mortality is implemented stochastically based on a general assumption of all-cause death of 10% of a population in a given year. Therefore, at the end of each day for each agent, there is a probability of a death occurring by randomly drawing a number and comparing it to 0.1/365, which represents the daily mortality risk equivalent to a 10% annual mortality risk. However, for generalists, to represent the fact that Matrix is the more risky habitat type according to their risk avoidance strategy, the mortality risk is increased to 13% if they are in a Matrix pixel. If a mortality event does occur, the specialist or generalist agent hatches a carcass agent which contains all of its attributes and is located on the same pixel, and then dies. The carcass is additionally assigned a value of day-died that matches the current value of the days-counter.

Nutrient transfer to pixels

Nutrient transfer from carcass agents to pixels is implemented through the check-patches submodel, which is run at the end of every day (every 8 timesteps). For carcasses where day-died has the same value as the current days-counter, then all nitrogen contained in the carcass (the body nitrogen, and any nitrogen consumed by the agent between the time that it last pooped and the time that it died) is transferred to the surrounding pixels. 50% of the total nitrogen is deposited in the pixel where the carcass is located, and the remaining 50% is distributed equally among the four nearest neighbors. This arrangement is meant to mimic the diffusion of nutrients by small scavengers (e.g. dung beetles: Veldhuis et al. 2017) and passive abiotic processes, although it does not account for longer-distance dispersal by highly mobile scavengers; white-backed vultures, for example, can forage over hundreds of kilometers (Phipps et al. 2013).

**Appendix II - Supplementary Figures & Tables**

**Table S1.** Simulation scenarios showing combinations of different population sizes and mean body size of specialists and generalists. Note that numbering of scenarios is not continuous because additional scenarios were run, but could not be used in direct comparisons to test our hypotheses and are thus not included here.

| Scenario Number | Mean Body Size (kg) | Number of Specialists | Number of Generalists |
|---|---|---|---|
| 1 | 10 | 0 | 50 |
| 2 | 10 | 0 | 100 |
| 3 | 10 | 0 | 150 |
| 4 | 10 | 0 | 200 |
| 5 | 25 | 0 | 40 |
| 6 | 25 | 0 | 50 |
| 7 | 25 | 0 | 60 |
| 8 | 25 | 0 | 80 |
| 9 | 25 | 0 | 100 |
| 10 | 25 | 0 | 150 |
| 11 | 40 | 0 | 25 |
| 12 | 40 | 0 | 37.5 |
| 13 | 40 | 0 | 50 |
| 14 | 40 | 0 | 100 |
| 15 | 40 | 0 | 150 |
| 31 | 10 | 50 | 0 |
| 32 | 10 | 100 | 0 |
| 33 | 10 | 150 | 0 |
| 34 | 10 | 200 | 0 |

| | | | |
|---|---|---|---|
| 35 | 25 | 40 | 0 |
| 36 | 25 | 50 | 0 |
| 37 | 25 | 60 | 0 |
| 38 | 25 | 80 | 0 |
| 39 | 25 | 100 | 0 |
| 40 | 25 | 150 | 0 |
| 41 | 40 | 25 | 0 |
| 42 | 40 | 37.5 | 0 |
| 43 | 40 | 50 | 0 |
| 44 | 40 | 100 | 0 |
| 45 | 40 | 150 | 0 |
| 61 | 10 | 400 | 0 |
| 62 | 25 | 160 | 0 |
| 63 | 10 | 0 | 400 |
| 64 | 25 | 0 | 160 |

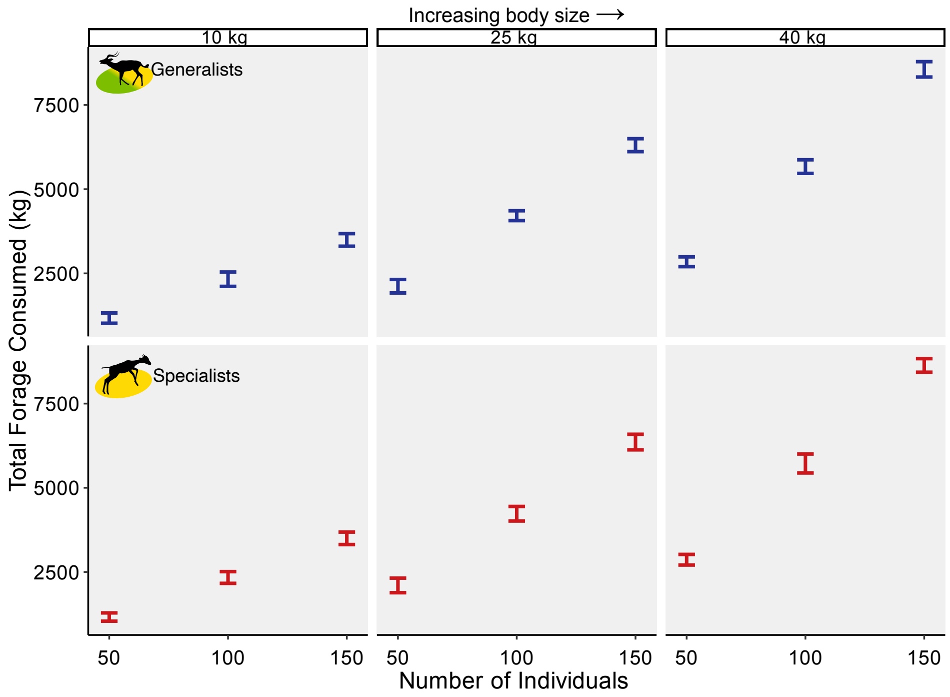


**Figure S1.** Relationship between the number of individuals and the total N consumed with population mean body mass of 10 kg, 25 kg, and 40 kg. Bars show mean value with standard deviation of 25 simulations for each scenario.

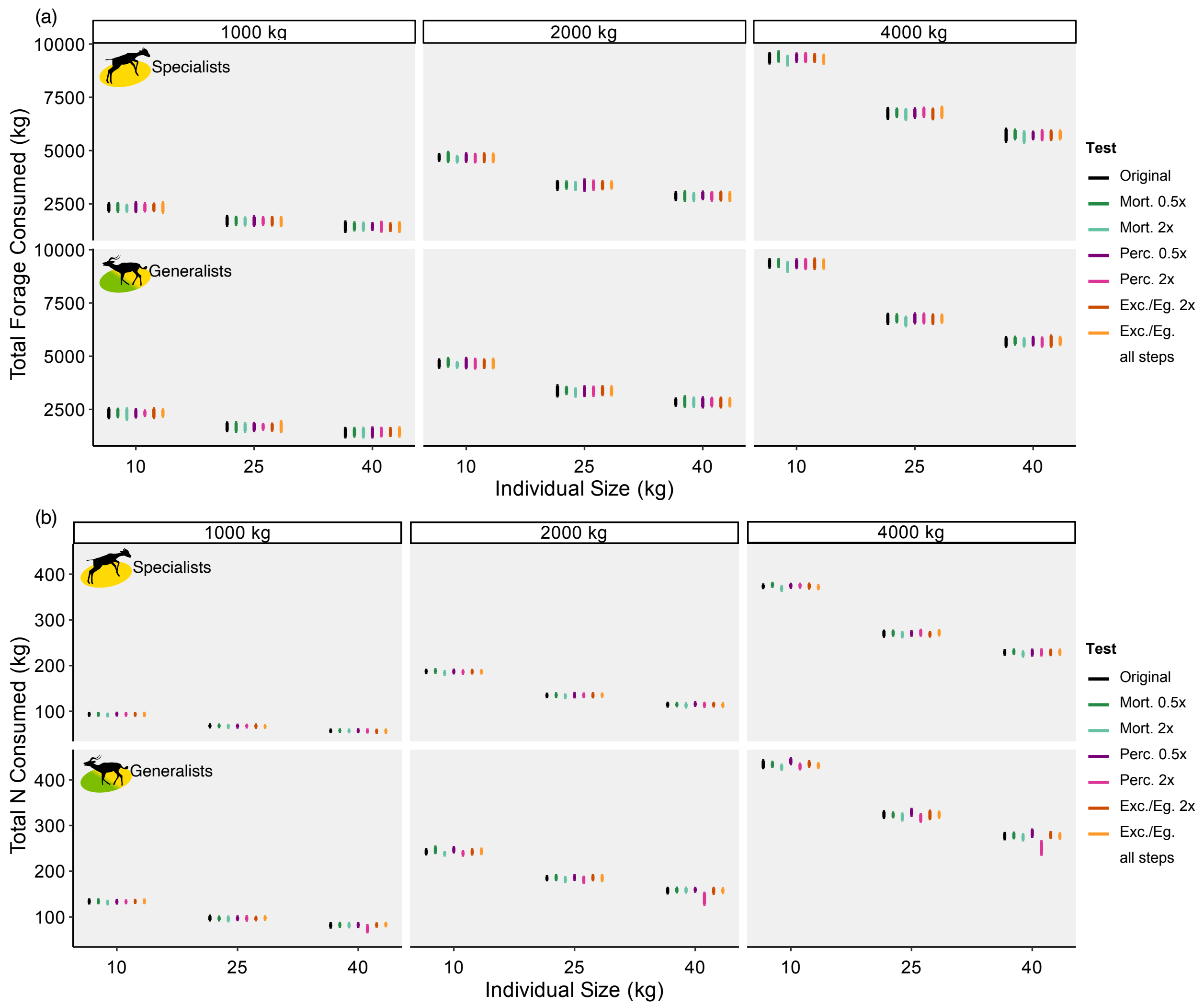


**Figure S2.** Relationship between the total biomass of individuals and (a) the total forage consumed, and (b) the total N consumed with population mean body mass of 10 kg, 25 kg, and 40 kg. Bars show mean value with standard deviation of 25 simulations for each scenario. Colors show sensitivity tests of key parameters, compared to the primary parameter set shown in **Figure 2** of the main text (indicated as "Original"): mortality rate either halved or doubled; perceptual range either halved or doubled; probability of excretion/egestion either doubled, or reaching 100% for every timestep.

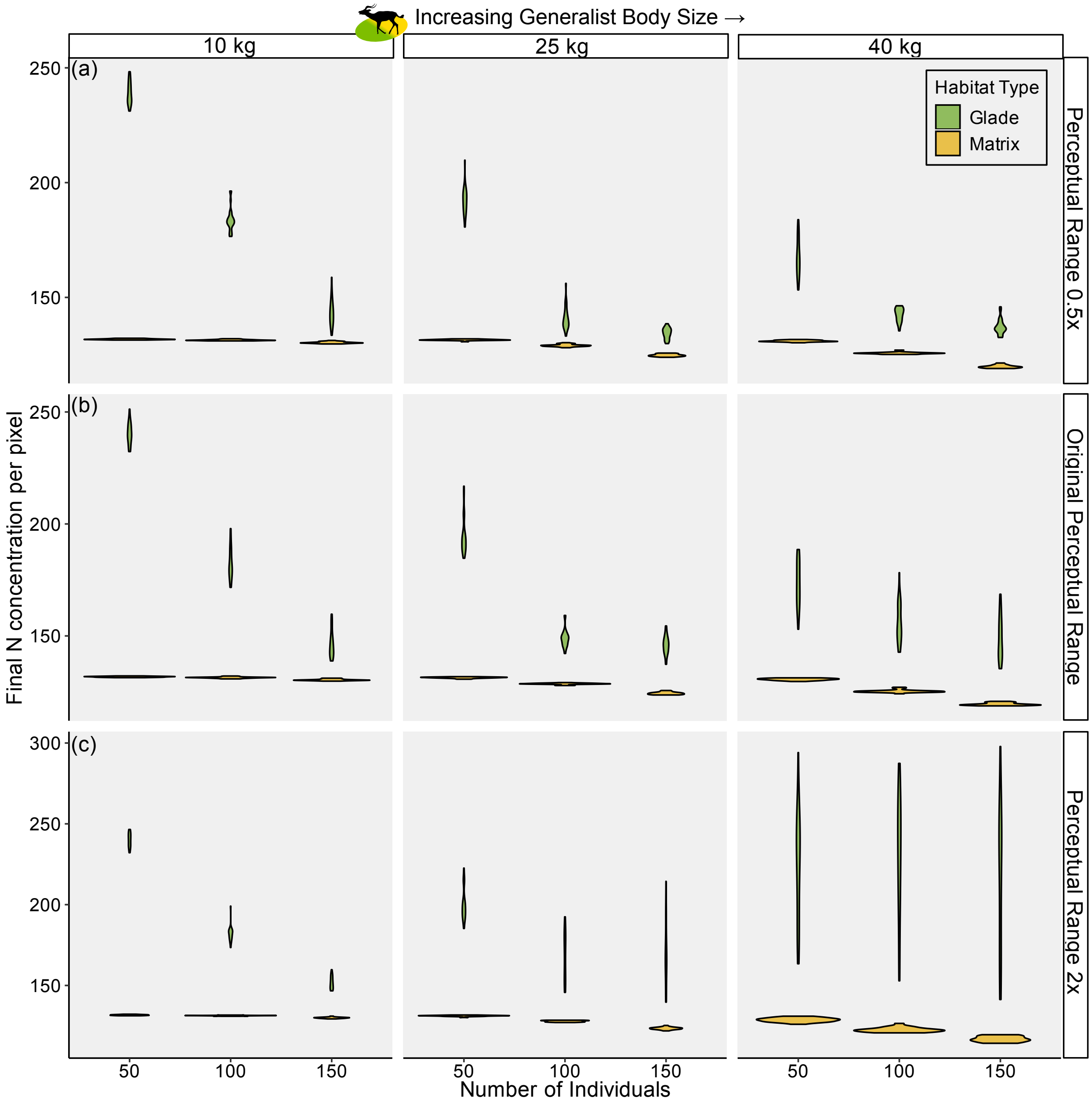


**Figure S3.** Average amount of nitrogen (in grams) contained in Glade (green) and Matrix (yellow) pixels at the end of simulations. Violin plots each show the distribution of data from 75 simulations. "Medium" perceptual range parameterization (shown in circles) corresponds to scenarios depicted in **Figure 5** in the main text. The "Lower" perceptual range is half this value, while the "Higher" perceptual range is twice this value. Note different y-axes in panels.

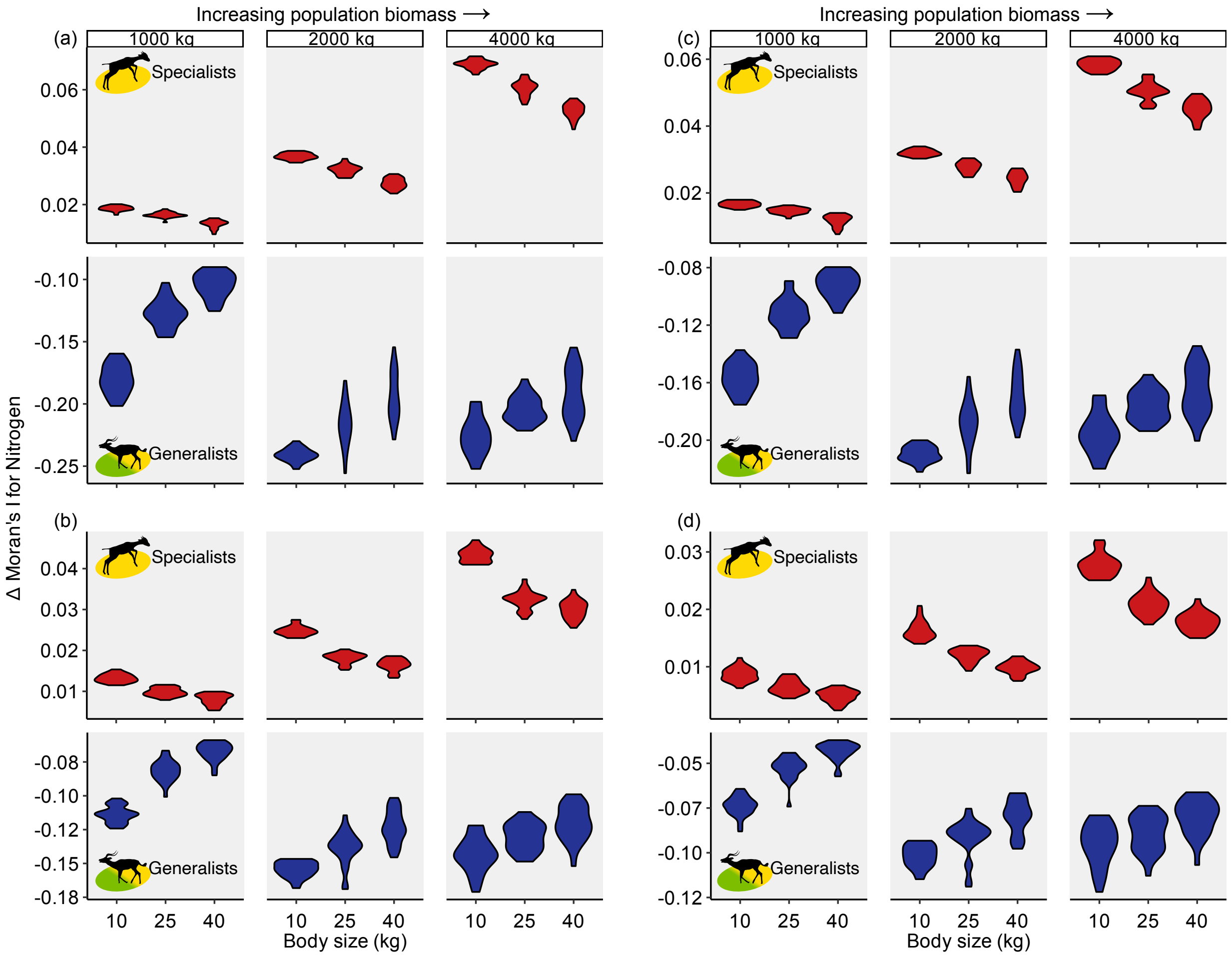


**Figure S4.** Change in Moran's I for the distribution of nitrogen between beginning and end of simulations when calculated over a neighborhood size of the 1 (a), 4 (c), 10 (b), or 17 (d) nearest neighbors. Plots show the effect of increasing mean body size (on the x-axis) when the total population biomass on the landscape is 1000 kg, 2000 kg, or 4000 kg of specialists (red violins) or generalists (blue violins). Violin plots show data from 25 replicate simulations for each scenario. Note different y-axes in panels.

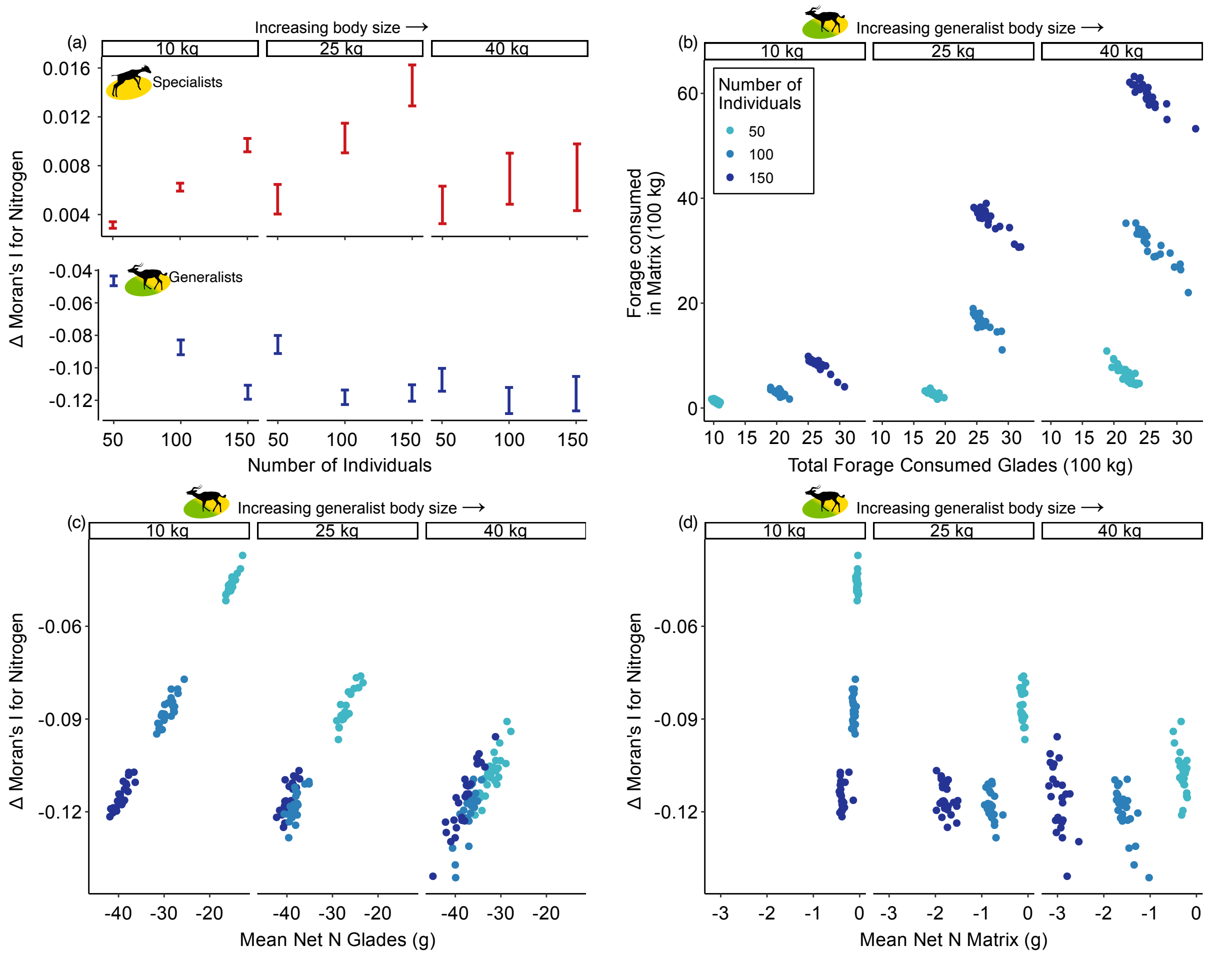


**Figure S5**. Key findings with the medium perceptual range level, but assuming that 75% rather than 50% of N consumed by herbivores is returned to the landscape via excretion and egestion, in simulations with mean individual body sizes of 10 kg, 25 kg, and 40 kg. In (a), bars show mean change in Moran's I and standard deviation of 25 simulations for each scenario. Note different y-axis scales for generalists and specialists; this panel allows direct comparison to **Figure S1**. (b) through (d) include only generalists, and colors differentiate simulations with 50, 100, and 150 individuals. (b) Relationship between total forage consumed in Glades and total forage consumed in Matrix. (c) and (d) Relationship between mean Net N in Glades (c) and Matrix (d), respectively, with the change in Moran's I. Panels (b), (c), and (d) allow direct comparison to **Figure 4**.

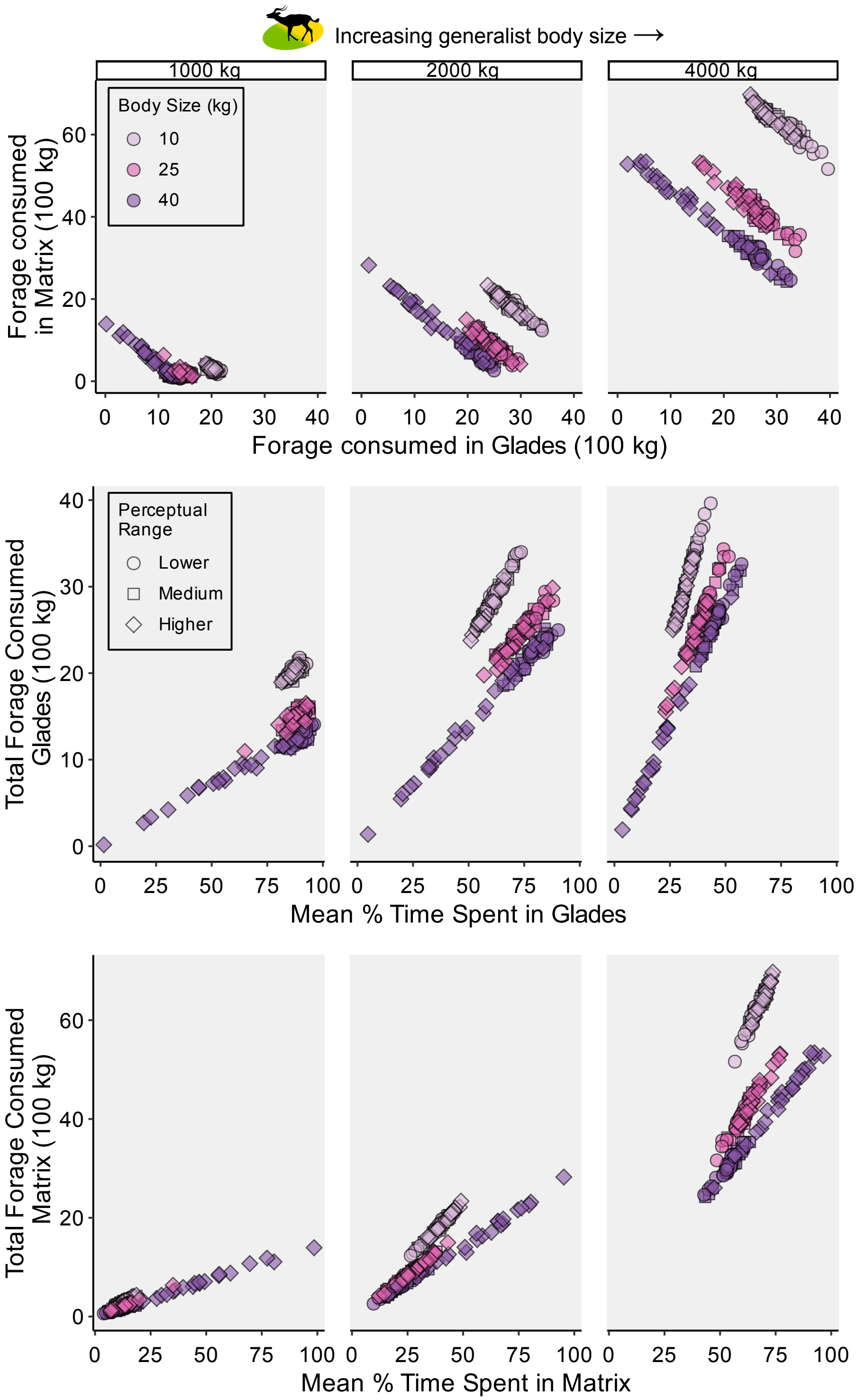


**Figure S6**. Effect of altering generalists' perceptual ranges on foraging patterns and nutrient distribution in the Glades and Matrix. The "Medium" perceptual range parameterization (shown in circles) corresponds to scenarios depicted in **Figure 5** in the main text. The "Lower" perceptual range is half this value, while the "Higher" perceptual range is twice this value. Apart from shapes on the graphs, all other figure elements are as in Figure 5.

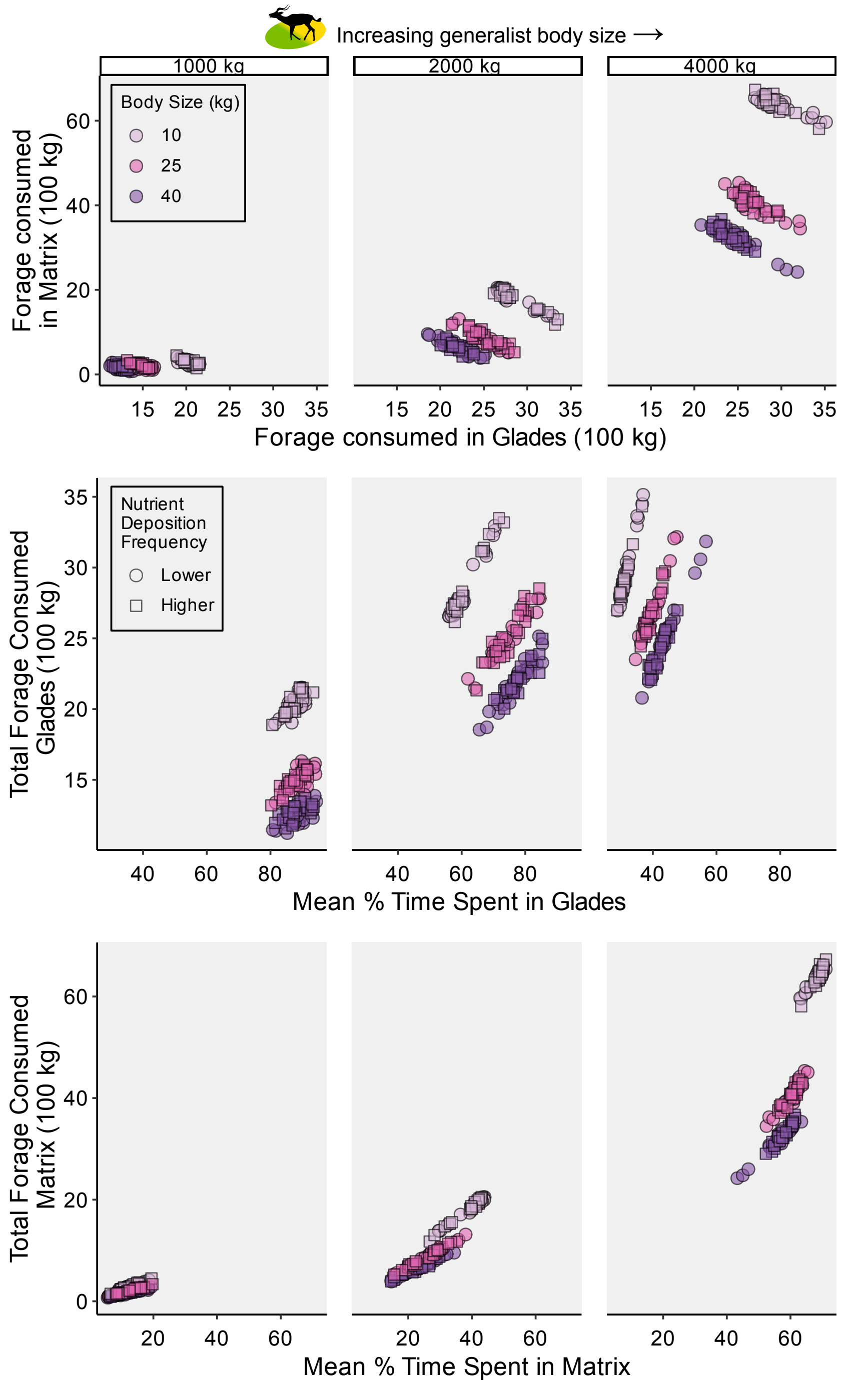


**Figure S7**. Effect of increasing the frequency of nutrient deposition by consumers, on foraging patterns and nutrient distribution in the Glades and Matrix. The "Lower" deposition parameterization (shown in circles) corresponds to the "Medium" scenarios depicted in **Figure 5** in the main text. The "Higher" deposition parameterization shows scenarios where consumers excrete and egest nutrients after every feeding bout, reducing the amount of nutrients in each deposition event but increasing the frequency and spatial proximity of depositions. Apart from shapes on the graphs, all other figure elements are as in Figure 5.

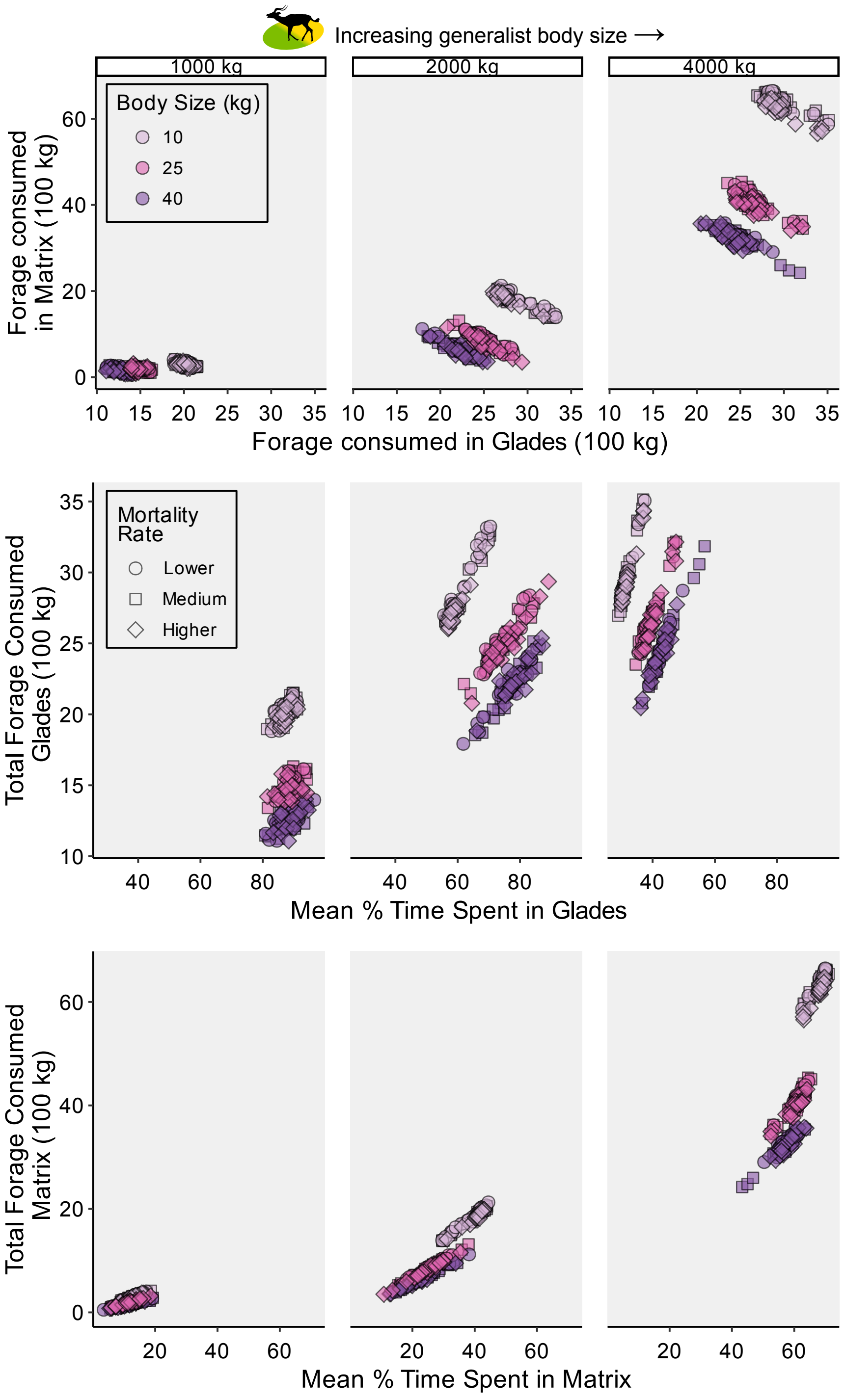


**Figure S8**. Effect of increasing the population-level mortality rate on foraging patterns and nutrient distribution in the Glades and Matrix. The "Medium" deposition parameterization (shown in squares) corresponds to the "Medium" scenarios depicted in **Figure 5** in the main text. The "Lower" parameterization (shown in circles) is with 50% of the original mortality rate, and the "Higher" parameterization (shown in diamonds) is with double the original mortality rate. Apart from shapes on the graphs, all other figure elements are as in Figure 5.
